# Single-nucleus multiome sequencing identifies candidate regulators of mouse gastric epithelial homeostasis

**DOI:** 10.64898/2026.04.23.720450

**Authors:** Maithê Rocha Monteiro de Barros, Katharina Bosch, Salima Soualhi, Shirin Issa Bhaloo, Thomas Chu, Tanya Hemrajani, Jin Cho, Kurtay Ozuner, Rui Fu, Heather Geiger, Nicolas Robine, Jade E.B. Carter, Silas Maniatis, Sandra Ryeom, Simon Tavaré, Karol Nowicki-Osuch

## Abstract

**Background & Aims:** Gastric epithelial cells maintain homeostasis through dynamic self-renewal mechanisms involving stem and progenitor cells; however, identifying them has been challenging. This study aims to identify stem cells of healthy gastric epithelium and cell type-specific regulators defining gastric epithelial homeostasis via single-nucleus multiome analysis.

**Methods:** Ten unique gastric samples were collected from 8-12 week old wildtype mice. Isolated nuclei were subjected to simultaneous profiling of gene expression and chromatin accessibility. After quality control, 31,598 cells were analyzed with Seurat and Signac using weighted-nearest neighbors analysis for joint RNA and ATAC clustering. Furthermore, SCENIC+, MultiVelo, EpiCHAOS and Cell plasticity score were used to uncover gene regulatory networks, cell state dynamics and lineage trajectories.

**Results:** Our analyses were validated by the identification of known regulators of stem-cell differentiation into mature cell types. More importantly, it revealed previously uncharacterized regulatory networks comprising novel transcription factor combinations that define cell identities, including *Ppara*, *Pparg*, *Arid5b* and *Sox5* as candidate regulators of parietal, foveolar, chief and neck cells, respectively. Further, our data support the identity of isthmus cells as stem-like cells of healthy gastric epithelium, as evidenced by epigenetic plasticity that simultaneously contains open chromatin states of all differentiated cell types in the absence of transcriptional reprogramming.

**Conclusion:** Consistent with Waddington’s epigenetic landscape hypothesis, gastric epithelial homeostasis is controlled by orchestrated epigenetic and transcriptional programs. Contrary to the prevailing hypothesis, stem cells can be defined not by a separate epigenetic state but by epigenetic superposition of differentiated cell states. Future work is needed to define the universality of these results.

## 1. Introduction

Human gastric epithelium is a columnar type of epithelium composed of single units called gastric glands, which are mosaics of multiple cell types including acid-secreting, digestive enzyme-secreting, mucous-producing and a plethora of hormone-secreting cells^1–4^. Gastric epithelium is organized in a monolayer and maintains homeostasis through self-renewal mechanisms involving stem and progenitor cells^1,5,6^. Human and mouse stomach contain corpus and antral glands featuring surface mucous foveolar cells, mucous neck cells, acid-secreting parietal cells, digestive enzyme-secreting chief cells, endocrine hormone-secreting cells and isthmus cells as the main epithelial cell types (Fig. 1a)^6^. Mouse gastric glandular epithelium is organized similar to human gastric glands. However, anatomical differences exist; for example, the murine forestomach is composed of squamous epithelium, akin to the esophageal squamous epithelium, a feature absent from human proximal stomach^7^.

**Figure 1.**
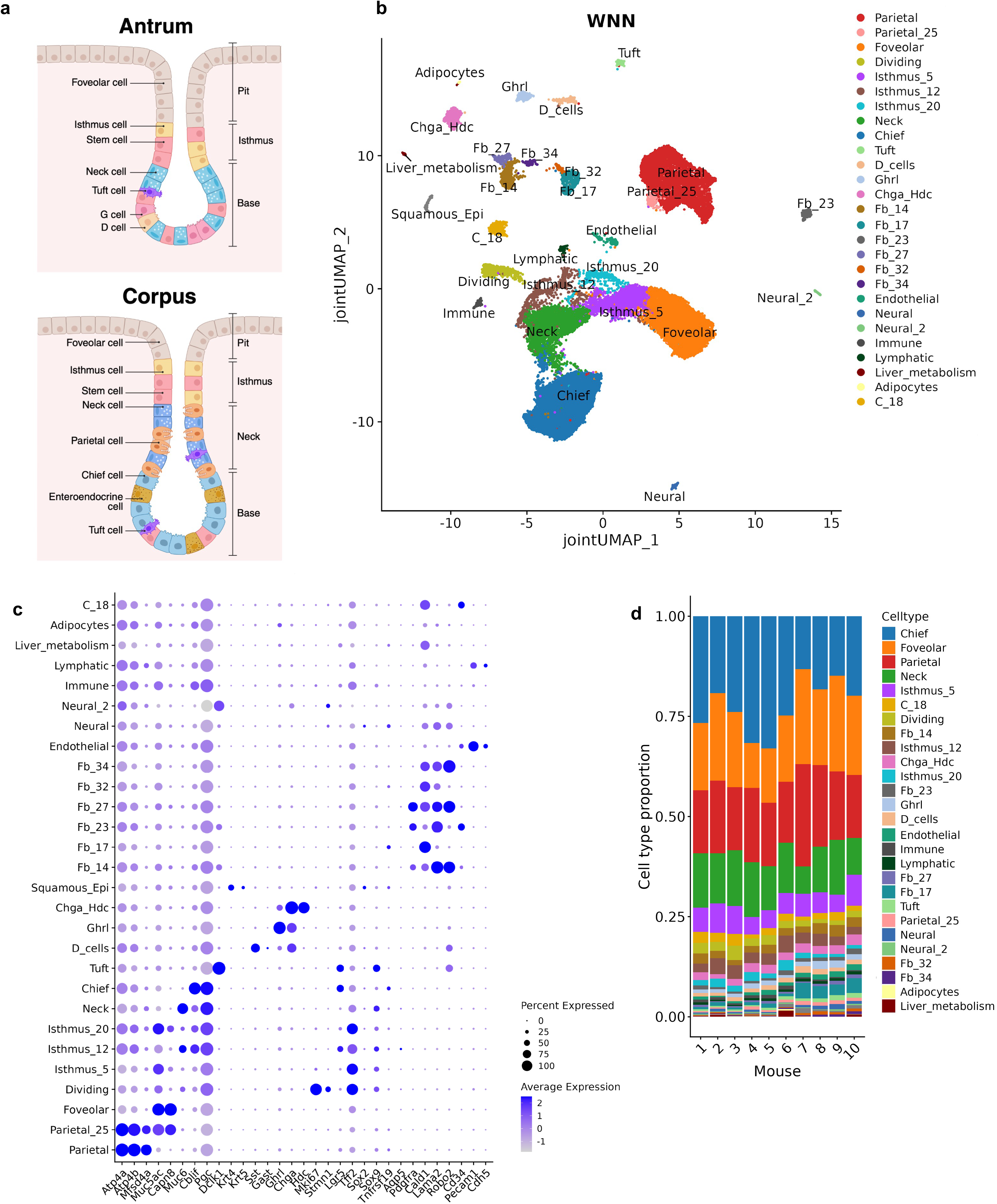
Integrated multiome profiling of the healthy mouse stomach successfully recovers all major epithelial and non-epithelial cell types. **a.** Diagram of main epithelial cell types found in the antrum and corpus regions of the stomach (Created with BioRender). **b.** WNN UMAP plot of 31,598 high-quality cells of integrated datasets after manual annotation featuring 27 different cell populations distinguished by transcriptional and epigenetic signatures. **c.** Dotplot showing expression of marker genes across cell populations. d. Stacked barplot displaying cell type distribution across each mouse.

Isthmus cells reside in the mid-gland area of gastric glands. During steady-state epithelial turnover, the isthmus region has been proposed as a source of stem cells and has been shown to replenish epithelial mature cell types bidirectionally^5,6,8^. However, it has also been shown that gastric chief cells can take on the role of stem cells and replenish the entirety of gastric epithelium, potentially in response to tissue injury ^9,10^.

Critical to the identity of a gastric stem cell is its ability to develop into all known gastric epithelial cell types. Over the years, numerous studies have investigated the identity of gastric stem cells using a variety of approaches, including organoid models and lineage tracing experiments. These studies span multiple regions of the stomach and focus on different stages of cellular differentiation. Candidate marker genes have been identified across these contexts, including *Lgr5*^11^, *Tff2*^12^, *Sox2*^13^, *Sox9*^14^, *Troy* (also known as *Tnfrsf19*)^15^, *Mist1*^16^ and *Aqp5*^17^, among others as discussed in various reviews^18–20^.

Despite these advances and seemingly well-known processes of gastric tissue self-renewal, there is a critical knowledge gap in the molecular mechanisms maintaining transcriptional and differentiation patterns of gastric epithelial cells. For example, only recently has the identity of a key transcription factor (TF) orchestrating the development of parietal cells been elucidated^21^. Crucially, the identity of gastric stem cells and the general mechanism of their self-renewal is poorly understood. Epigenetic reprogramming might be the key factor controlling gastric cell lineages and the differentiation of gastric epithelium might follow the epigenetic landscape hypothesis proposed by Waddington^22^ in 1957. Modern interpretations of this concept suggest that cellular states are probabilistic and individual cell types can have some capacity to develop into other cell types by acquisition of alternative cellular states, which is defined as cellular plasticity^23^. However, direct observation of such states outside of hematopoietic differentiation is limited.

Knowledge of gastric stemness with a detailed understanding of how cell lineages are specified and maintained under homeostatic conditions, is fundamental to revealing the mechanisms that drive gastric cancer (GC) and for identifying its cells of origin. GC ranks fifth worldwide in both incidence and mortality, with 968,784 cases and 660,175 deaths in 2022^24^. It is a complex multi-factorial, multi-step heterogeneous disease that can follow sequential events described as Correa’s Cascade^25,26^. This cascade highlights progressive morphological and phenotypical changes that take place in the gastric epithelium, starting at the normal gastric mucosa, going through non-atrophic gastritis, chronic atrophic gastritis, intestinal metaplasia, low grade dysplasia, high grade dysplasia and finally invasive adenocarcinoma. At the heart of Correa’s Cascade lies gradual loss of normal gastric homeostasis, often in response to *Helicobacter pylori* infection.

Recent advances in sequencing technologies have opened new avenues for studies of gastric homeostasis in unparalleled detail. In this study we characterize the transcriptional and epigenetic landscape of healthy mouse stomach tissue using paired single-nuclei RNA (snRNA-seq) and ATAC sequencing (snATAC-seq). Using gene expression and accessible chromatin analysis, we annotated 27 major cell type populations, including the identification of a new population of cells (Parietal_25 cluster) that display epigenetic features of both parietal and foveolar cells, indicating that these two cell types share lineage trajectories. Through enhancer-driven gene regulatory network (eGRN) and various trajectory analyses, we identified regulatory programs driving gastric epithelium cellular states with a particular focus on stemness and cell identity. Our approach identified previously described regulators of differentiated cell types such as *Esrrg* for parietal cells^27^, *Klf4* for foveolar cells^28^, *Bhlha15* (also known as *Mist1)* for chief cells^16^ and *Sox9* for neck cells^29^. Furthermore, we identified *Ppara, Pparg, Arid5b* and *Sox5* as new putative regulators in parietal, foveolar, chief and neck cells respectively. We also identified three isthmus cell clusters that are highly plastic (Isthmus_5, Isthmus_12, Isthmus_20), lacking specific stem-like markers, but are epigenetically primed to become mature epithelial stomach cell types. Furthermore, in silico TF perturbation analysis supported the functional relevance of the predicted regulators governing epithelial cell identity.

## 2. Results

### Multiomics of healthy gastric tissue uncovers epithelial cell identity and reveals stem-like populations

To characterize gastric cell populations in mice, we used 10x Genomics single cell Multiome (scMultiome) to simultaneously profile gene expression (snRNA-seq) and chromatin accessibility (snATAC-seq) in nuclei isolated from the gastric columnar tissues of 10 healthy mice, with samples processed in two sequencing batches of six and four mice (Supplementary Table 1). After quality control, a total of 31,598 nuclei were analyzed using the Seurat^30^ and Signac packages^31^. Weighted-nearest-neighbors (WNN) analysis allowed for joint clustering of the RNA and ATAC assays^32^. We identified a total of 24,596 genes and 514,311 regulatory regions.

Unsupervised clustering of the WNN assay resulted in the identification of 38 clusters (Supplementary Fig. 1a), which were manually annotated based on known cell type markers^33–35^ and cluster-specific markers after differential gene expression analysis. Following annotation, the clusters were merged into 27 major cell populations (Fig. 1b,c). All clusters were composed of cells from all biological replicates in comparable proportions, with chief, foveolar, parietal and neck representing the most abundant populations (Fig. 1d).

Epithelial populations were composed of foveolar (*Muc5ac*), neck (*Muc6)*, parietal (*Atp4a, Atp4b, Esrrg*), chief (*Cblif, Pgc*) and dividing (*Mki67*) cells. Additionally, we identified all known gastric enteroendocrine cell types including D and G cells mixed in a cluster (*Sst, Gast –* “D_cells”), Tuft cells (*Dclk1*), P/D1 or X/A-like cells (*Ghrl*) and enterochromaffin-like (*Chga*, *Hdc*) cells and a small cluster of squamous cells (*Krt4, Krt5 –* “Squamous_Epi”). Non-epithelial clusters were composed of endothelial, immune, lymphatic and neural cells, as well as six different fibroblast clusters (*“Fb”* followed by the original unsupervised cluster number), adipocytes, a cluster linked to liver (“Liver_metabolism”) and a cluster of unknown origin (“C_18”).

Three clusters did not align with any established categories of fully differentiated gastric cells. Instead, they expressed markers associated with isthmus and stem-like cells such as *Lgr5, Tnfrsf19* and *Tff2* alongside other well-characterized gastric markers such as *Atp4a, Muc5ac, Muc6, Cblif* and *Pgc* (Fig. 1c). As a result, these clusters were named Isthmus_5, Isthmus_12, Isthmus_20 according to the original unsupervised cluster from which they emerged. Similarly, another cluster expressed markers of both parietal (*Atp4a, Atp4b, Mfsd4a)* and foveolar (*Muc5ac, Capn8*) cells and was named Parietal_25.

Next, we employed trajectory analysis tools and took advantage of the multimodal nature of our data to uncover differentiation patterns of epithelial cells (clusters Parietal, Parietal_25, Foveolar, Dividing, Isthmus_5, 12 and 20, Neck, Chief, Tuft, D cells, Ghrl and Chga_Hdc) (Fig. 2a). Firstly, we used MultiVelo^36^ to characterize the latent time developmental pattern of gastric epithelium (Fig. 2c,d). We then used previously described measures of discordance between gene expression and chromatin states to define cellular plasticity^37^ (Fig. 2e). Finally, we employed the EpiCHAOS score^38^ to measure within cluster epigenetic diversity derived from the scATAC-seq data (Fig. 2f).

**Figure 2.**
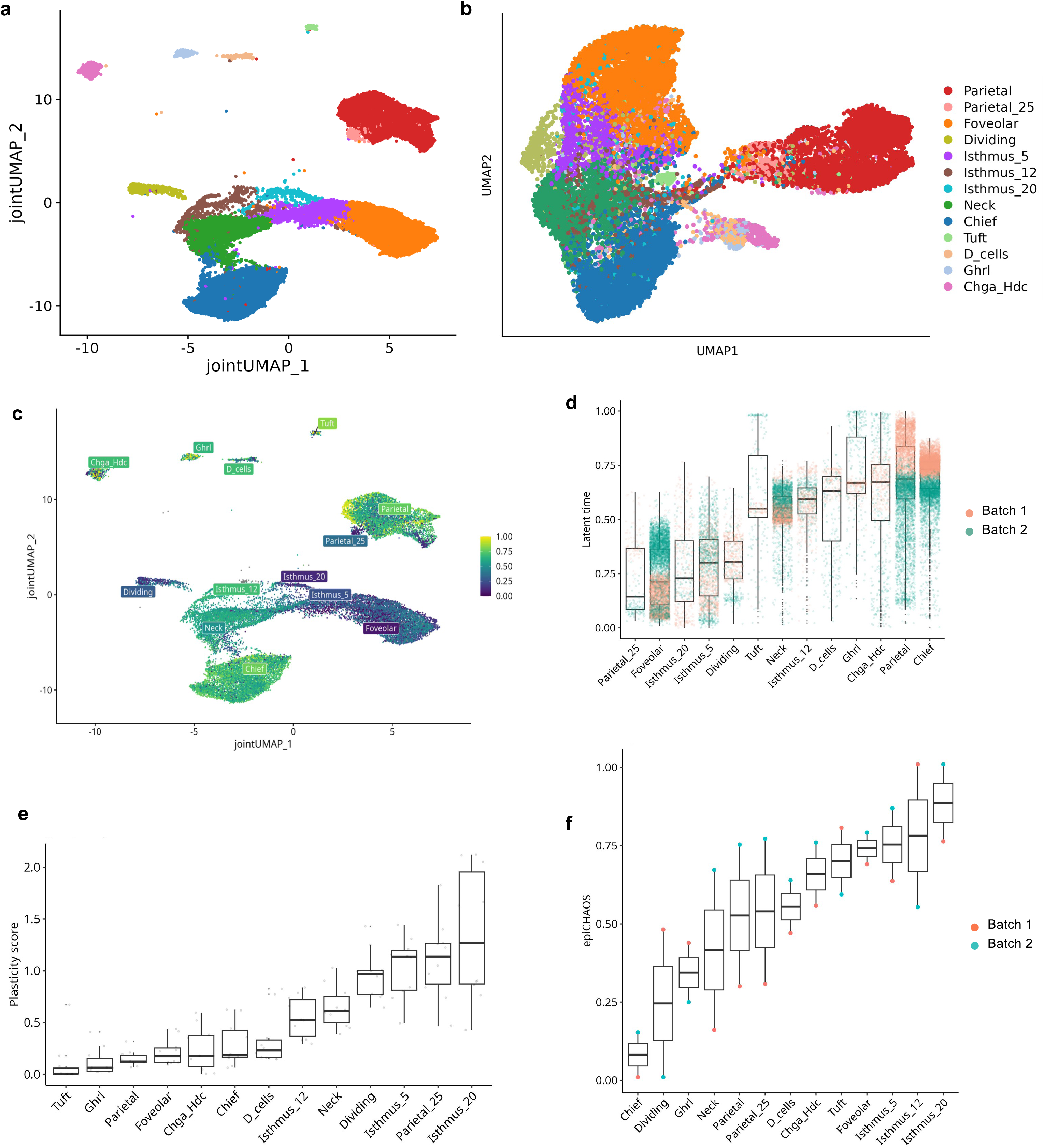
SCENIC+ and trajectory analyses reveal dynamic states and plasticity of epithelial cell types. **a.** Seurat Multiome WNN UMAP plot of 27,810 cells featuring 13 epithelial clusters distinguished by transcriptional and epigenetic signatures. **b.** SCENIC+ UMAP plot of epithelial cells based on eRegulon target gene and target region enrichment scores, representing cell type clustering based on active regulatory networks. **c.** MultiVelo latent time projected on the WNN embedding representing the predicted developmental progress of individual cells, inferred from the temporal relationship between chromatin accessibility and gene expression. Higher values identify more differentiated cell states. **d.** Distribution of MultiVelo latent time displayed as boxplots for both datasets, reflecting the predicted developmental progress of cell populations. **e.** Metacell plasticity score boxplot across epithelial cell types with dots representing individual animals, quantifying the uncertainty of cell state predictions based on chromatin accessibility profiles. High scores indicate cells whose epigenome is compatible with multiple potential transcriptional identities **f.** Boxplot displaying EpiCHAOS scores across epithelial cell types for both datasets. Higher values indicate greater within-cluster cell-to-cell epigenetic heterogeneity.

We also used SCENIC+ to identify eGRNs for the epithelial cell types of interest by mapping each TF’s target regions and target genes, thereby inferring enhancer-driven regulatory relationships (eRegulons) (Fig. 2b). We built an atlas of gastric epithelium eRegulons composed of direct and extended categories. A total of 468 eRegulons were identified, 170 direct and 298 extended, with an average of 407 target regions and 137 target genes and 478 target regions and 147 target genes, respectively. After further filtering (Methods), and in line with the guidelines of SCENIC+, only positive region-to-gene (R2G) eRegulons were kept^39^. As a result, 160 eRegulons (94 direct, 66 extended) were used in the downstream analysis. Regulon specificity score (RSS) (Fig. 3a) and area under the curve (AUC) (Fig. 3b) values were used to link eRegulons to cell types. SCENIC+ not only recovered known regulators for parietal, foveolar, neck and chief cells, but also uncovered new potential regulators and highlighted candidate enhancer regions that show cell type specific accessibility for most cell types in our dataset.

**Figure 3.**
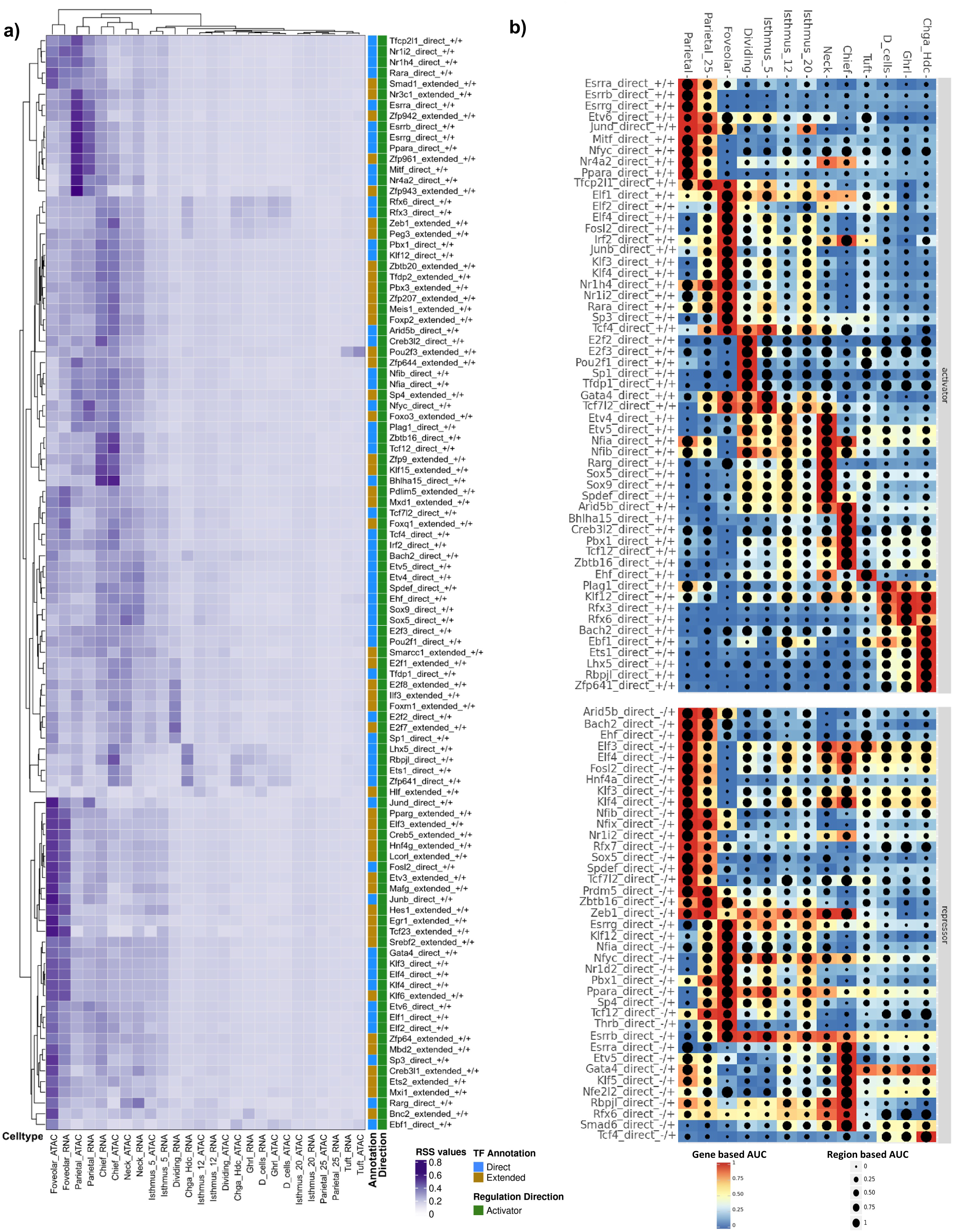
SCENIC+ cell specificity scores (RSS) and eRegulon AUC enrichment scores reveal regulatory programs associated with epithelial cell identities. **a.** Heatmap of Regulon cell specificity scores (RSS) for activator eRegulons across cell types based on the RNA and ATAC assays. Transcription factor (TF) annotation can be direct (blue) or extended (yellow). RSS values are calculated via Jensen-Shannon divergence to quantify the lineage-specific activity of each eRegulon. Heatmap is clustered by cell type (columns) and eRegulons (rows). **b.** Heatmap-dotplot of AUC gene (color) and AUC region (dot size) values per activator and repressor direct eRegulons across cell types. AUC values measure the enrichment of a regulon’s target nodes within a cell’s overall ranking of gene expression and chromatin accessibility. Cell types are ordered by transcriptomic similarity to highlight the dynamic shifts in regulatory networks.

### Gene regulatory network analysis uncovers putative regulators of foveolar and parietal differentiation

UMAP projection based on the contribution of each eRegulon to individual cell types captured three major cellular patterns of differentiation. Unlike the WNN-based projections (Fig. 2a), parietal cells formed via a continuous differentiation pattern from the isthmus cell types via parietal_25 cells (Fig. 2b). We further observed a foveolar differentiation path and a secretory path via neck and chief cells as previously seen in mice^40^. Various enteroendocrine cell types formed joint differentiation paths linked to the isthmus and chief cell types. MultiVelo analysis also showed that parietal cells display one of the highest latent time values (Fig. 2c,d) and one of the lowest plasticity scores across epithelial cell types (Fig. 2e), showing that this cell type is highly differentiated and specialized.

Cluster parietal_25 showed features of putative parietal cell progenitor population. Cells within that cluster express key markers of parietal cells such as *Atp4a, Atp4b* and *Mfsd4a* (Fig. 1c). However, we observed a critical lack of synchronization between the RNA and ATAC-seq states as captured by UMAP dimensionality reduction. The RNA state indicates similarity between parietal_25 and mature parietal cell types. However, the ATAC modality places them within foveolar and isthmus_5 cellular states (Fig. 4a,b,c). Closer examination of the open chromatin states in the target regions of key parietal (*Esrrg*) and foveolar (*Klf4*) eGRNs TFs indicated that these cells simultaneously possess epigenetic features of both cell types (Fig. 4d,e). Thus, our data indicate that parietal and foveolar cells share lineage trajectories with a split occurring shortly after differentiation from the isthmus cell types with potential early expression of *Esrrg* as a key parietal lineage commitment marker.

**Figure 4.**
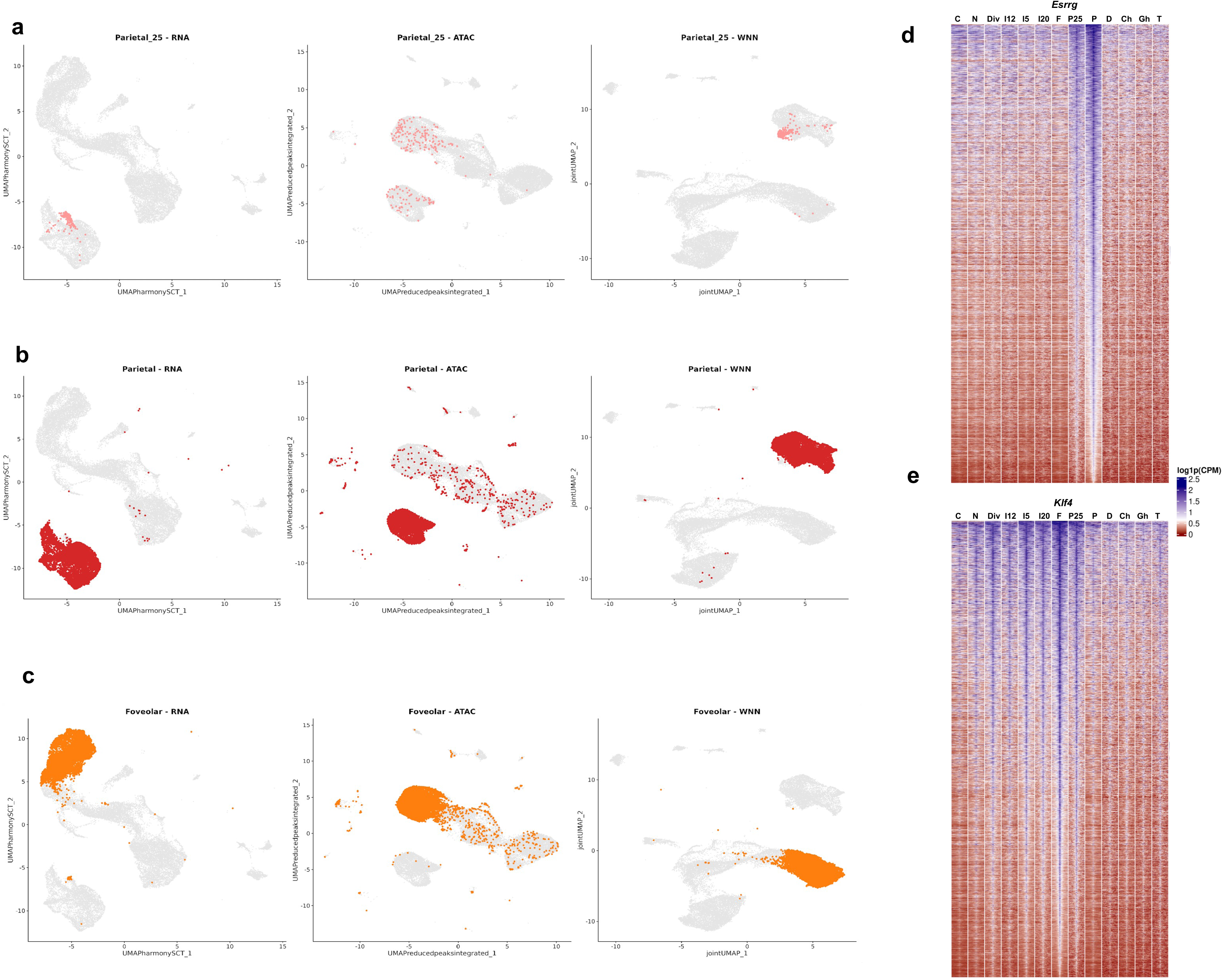
Parietal_25 cluster shows epigenetic features of parietal and foveolar cell types. RNA, ATAC and WNN UMAP embeddings highlighting the distribution of **a.** Parietal_25, **b.** Parietal and **c.** Foveolar cells demonstrating their transcriptional, chromatin-accessibility and integrated multimodal identities. Heatmap showing chromatin accessibility across regions assigned to the **d.** Esrrg direct activator eRegulon linked to Parietal cells and **e.** Klf4 direct activator eRegulon linked to Foveolar cells. C:Chief, N:Neck, Div:Dividing, I5:Isthmus_5, I12: Isthmus_12, I20:Isthmus_20, F: Foveolar, P25:Parietal_25, P:Parietal, D:D_cells, Ch:Chga_Hdc, Gh:Ghrl, T:Tuft cells.

*Esrrg*, estrogen related receptor gamma, is a recently discovered regulator in the differentiation and maturation of stem cells into acid-secreting parietal cells^27^. In line with its role, *Esrrg* is highly expressed in parietal cells compared to the other epithelial cell types, and the expression of its target genes follows the same pattern (Fig. 5a). Detailed investigation of the top 50 target genes based on TF-to-Gene (TF2G) importance demonstrated that they are highly and selectively expressed in parietal cells with lower expression in the parietal_25 cluster (Fig. 5b). Chromatin accessibility around binding regions for all TFs in the eGRN was also highly specific to parietal cells with some overlap in parietal_25 cells (Fig. 5c). Our identification of *Esrrg* serves to validate our computational approach. The analysis uncovered four further candidate regulators driving cell fate decisions in parietal cells: *Esrra, Esrrb*, *Ppara* and *Mitf* (Fig. 5d). Triple perturbation of *Esrra, Esrrb* and *Esrrg* highlighted their cooperative role in maintaining parietal identity (Supplementary Fig. 4a).

**Figure 5.**
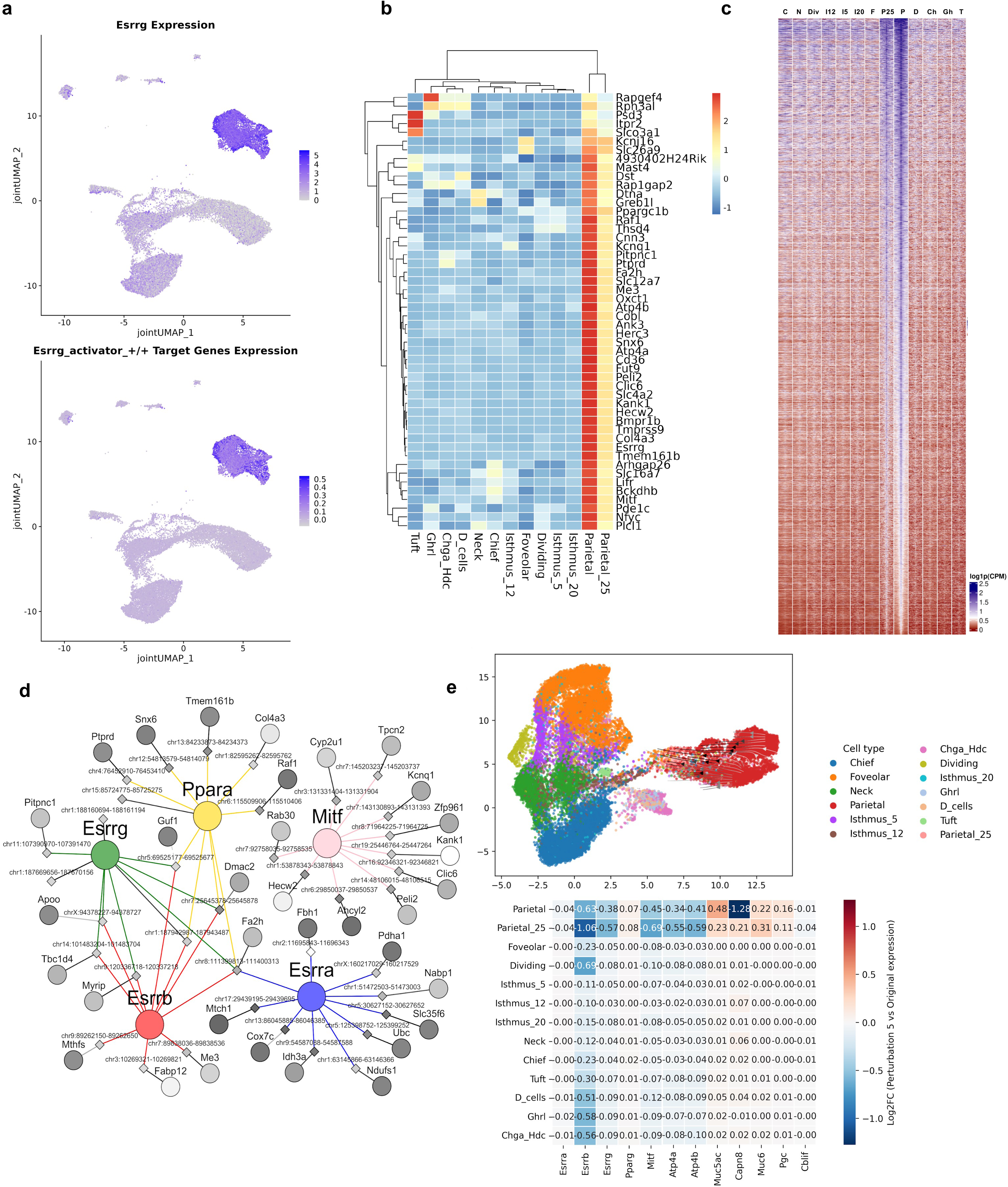
Parietal cell regulatory program recovering the role of Esrrg and nominating Ppara as a novel regulator. **a.** UMAP displaying Esrrg expression and the expression of its target genes across epithelial clusters. **b.** Heatmap showing expression of top 50 target genes based on TF-to-gene importance values in the Esrrg direct activator eRegulon. **c.** Heatmap showing accessibility of all regulatory regions in the parietal eGRN across cell types. **d.** Visualization of the eGRN of parietal cells limited to the top 10 edges per TF based on triplet ranking values. Colorful circles represent TFs, diamonds represent enhancer regions and smaller circles represent target genes. Region-to-gene edges are shaded by their importance scores (importance_R2G) and nodes are shaded by log2FC values of expression and chromatin accessibility, where darker shades indicate higher values. **e.** Visualization of the effect of Ppara in silico perturbation simulation after 5 iterations. Projected effect in the SCENIC+ UMAP embedding where arrows indicate each cell’s shift in embedding space, shaded by distance traveled, with darker arrows marking larger movements. Heatmap displaying log2FC between original gene expression and perturbed expression across epithelial cell types and genes of interest.

In addition to the previously described Esrr family of TFs, we also identified two novel putative regulators of parietal cells: *Ppara* (Peroxisome proliferator-activated receptor alpha) and *Mitf* (Microphthalmia-associated transcription factor). Previous studies linked *Mitf* with the differentiation of isthmus cells into parietal cells^40^, but no role for *Ppara* has been previously reported. In silico knockdown of *Ppara* generated a pronounced shift in parietal cells towards parietal_25, as demonstrated by the co-embedding of the perturbed gene expression matrix with the original one. Similarly, this perturbation led to a downregulation of all TFs in the parietal eGRN (*Esrra, Esrrb, Esrrg, Mitf*) and of known parietal markers (*Atp4a, Atp4b*) but led to an upregulation of foveolar marker genes (*Muc5ac*) in both parietal and parietal_25 cells (Fig. 5e).

On the other hand, simulated knockdown of *Pparg* produced a predicted transition from foveolar cells towards parietal_25, dividing and isthmus_5 with an upregulation of most parietal eGRN TFs (*Esrrb, Esrrg, Ppara*) and marker genes (*Atp4a, Atp4b*) in foveolar cells and an overall downregulation of foveolar markers (*Muc5ac, Capn8*) across all cell types (Supplementary Fig. 3a). The comparative analysis of *Ppara* (Fig. 5e) and *Pparg* (Supplementary Fig. 3a) demonstrated that they have mutually exclusive expression and downstream eRegulon activation patterns. Since *Ppara* promotes lipid catabolism and *Pparg* lipid storage^41,42^, their opposite functions might indicate that metabolic differentiation is required for appropriate separation of foveolar and parietal cell lineages.

In addition to *Pparg,* SCENIC+ predicted cooperativity between *Sp3*, *Elf2*, *Elf4*, *Klf3*, *Klf4*, *Fosl2*, *Junb* and *Rara* in the regulation of foveolar cells, as demonstrated by the eGRN (Fig. 6d). Only the role of *Klf4* has previously been linked with gastric epithelial proliferation and differentiation towards mucous cell populations^43^. *Klf4* and its target genes are highly expressed in foveolar cells compared to the other cell types (Fig. 6a). These target genes also showed moderate expression in isthmus clusters (Isthmus_5 and Isthmus_20), parietal_25 cluster as well as the dividing cluster (Fig. 6b). The accessibility at all target regulatory regions was specific to foveolar cells with some regions open in parietal_25, dividing, isthmus_5 and isthmus_20 clusters (Fig. 6c).

**Figure 6.**
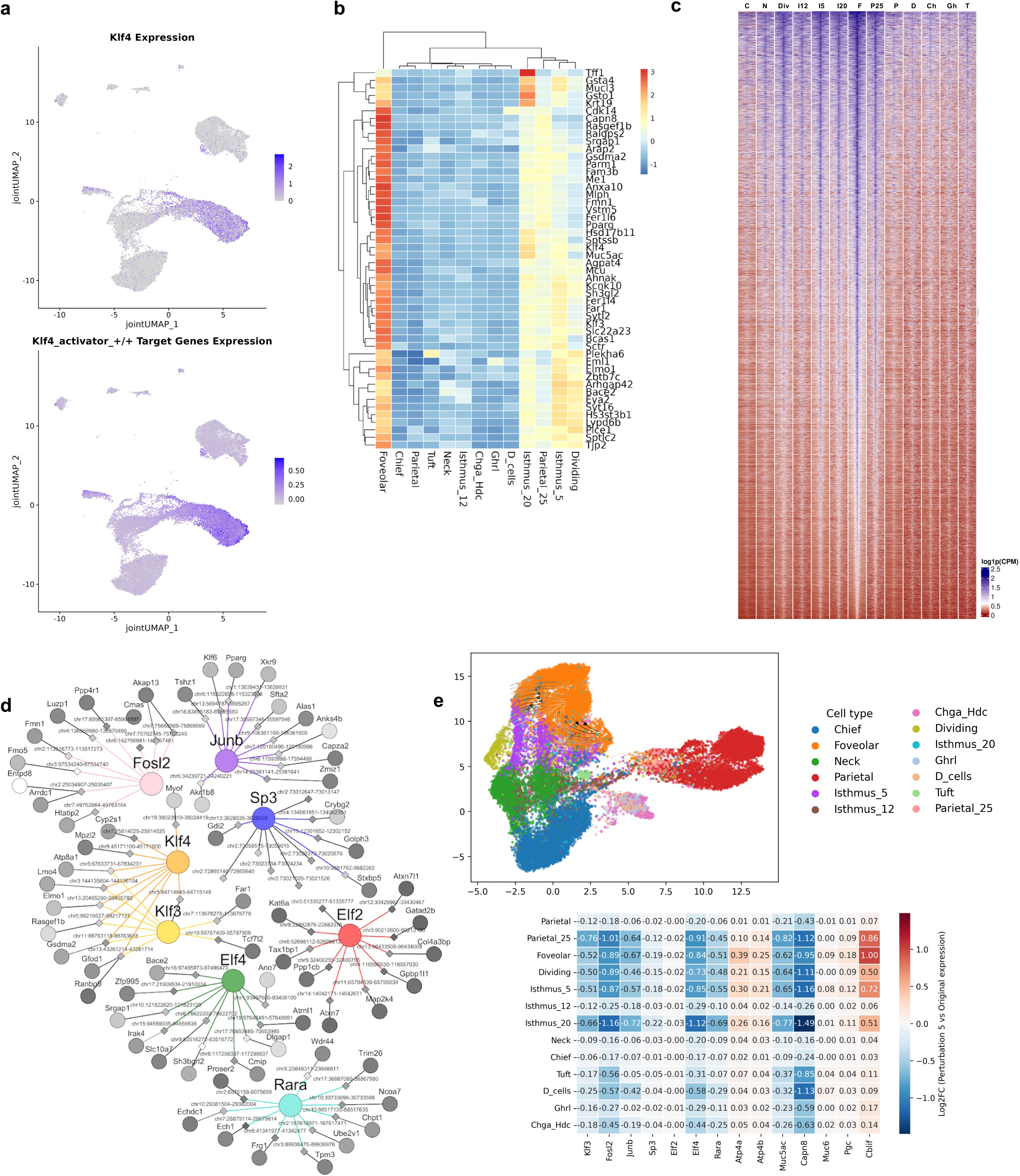
Foveolar cell regulatory program recovering the role of Klf4 as a master regulator. **a.** UMAP displaying Klf4 expression and the expression of its target genes across epithelial clusters. **b.** Heatmap showing expression of top 50 target genes based on TF-to-gene importance values in the Klf4 direct activator eRegulon. **c.** Heatmap showing accessibility of all regulatory regions in the foveolar eGRN across cell types. **d.** Visualization of the eGRN of foveolar cells limited to the top 10 edges per TF based on triplet ranking values. Colorful circles represent TFs, diamonds represent enhancer regions and smaller circles represent target genes. Region-to-gene edges are shaded by their importance scores (importance_R2G) and nodes are shaded by log2FC values of expression and chromatin accessibility, where darker shades indicate higher values. **e.** Visualization of the effect of Klf4 in silico perturbation simulation after 5 iterations. Projected effect in the SCENIC+ UMAP embedding where arrows indicate each cell’s shift in embedding space, shaded by distance traveled, with darker arrows marking larger movements. Heatmap displaying log2FC between original gene expression and perturbed expression across epithelial cell types and genes of interest.

In silico knockdown of *Klf4* led to a global downregulation of TFs involved in the foveolar eGRN (*Klf3*, *Fosl2, Junb, Sp3, Elf2*, *Elf4*, *Rara*) and foveolar gene markers (*Muc5ac, Capn8*) across all cell types but mainly in parietal_25, foveolar, dividing, isthmus_5 and isthmus_20, confirming that *Klf4* is necessary to maintain the foveolar phenotype (Fig. 6e). Interestingly, this perturbation also led to an upregulation of chief cell marker *Cblif* across all cell types, but mainly in parietal_25, foveolar and isthmus_5.

### Neck and chief cells share differentiation trajectory

Next, we turned our attention to the secretory cell types. SCENIC+ recovered a known master regulator of chief cells, the TF *Bhlha15*, also known as *Mist1*^16^. The investigation of expression patterns and target regulatory regions demonstrated the highly specific role of *Bhlha15* (Fig. 7a, b) with potential interactions with *Arid5b* and *Zbtb16* TFs (Fig. 7d). Chromatin accessibility at the binding sites of *Bhlha15, Arid5b, Creb3l2, Pbx1, Tcf12* and *Zbtb16* within the eGRN was highly specific to chief cells with a partial overlap with isthmus_12 and isthmus_20 clusters. It also showed that some of these chromatin states are already present in the neck cells, suggesting differentiation from the neck to chief cell types (Fig. 7c).

**Figure 7.**
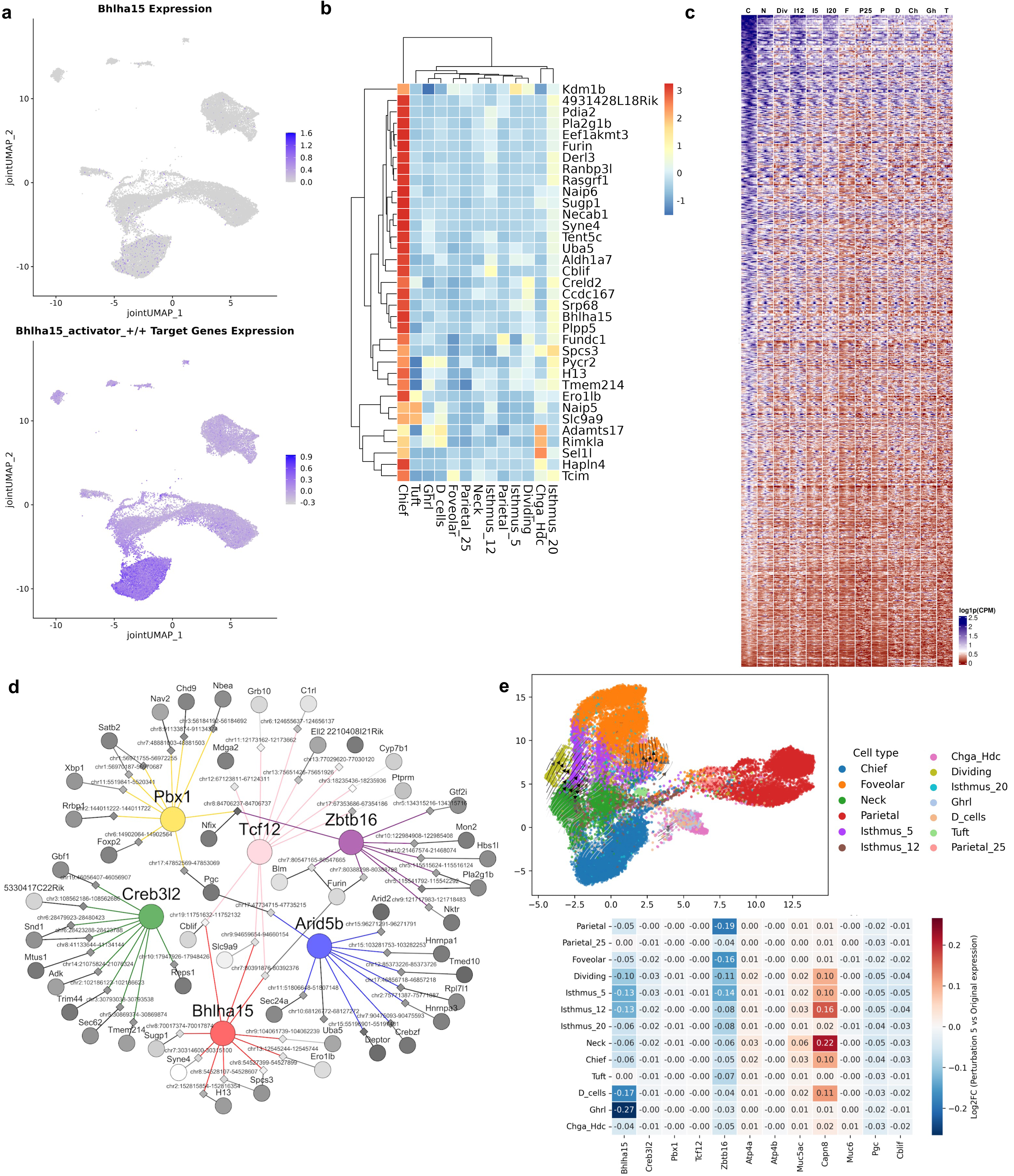
Chief cell regulatory program recovering the role of Bhlha15 and suggesting Arid5b as a novel regulator. **a.** UMAP displaying Bhlha15 expression and the expression of its target genes across epithelial clusters. **b.** Heatmap showing expression of top 50 target genes based on TF-to-gene importance values in the Bhlha15 direct activator eRegulon. **c.** Heatmap showing accessibility of all regulatory regions in the chief eGRN across cell types. **d.** Visualization of the eGRN of chief cells limited to the top 10 edges per TF based on triplet ranking values. Colorful circles represent TFs, diamonds represent enhancer regions and smaller circles represent target genes. Region-to-gene edges are shaded by their importance scores (importance_R2G) and nodes are shaded by log2FC values of expression and chromatin accessibility, where darker shades indicate higher values. **e.** Visualization of the effect of Arid5b in silico perturbation simulation after 5 iterations. Projected effect in the SCENIC+ UMAP embedding where arrows indicate each cell’s shift in embedding space, shaded by distance traveled, with darker arrows marking larger movements. Heatmap displaying log2FC between original gene expression and perturbed expression across epithelial cell types and genes of interest.

We recovered candidate enhancers linking *Bhlha15* with the main markers of chief cells: *chr19:11751632-11752132* for *Cblif*, and *Arid5b* with *chr17:47734715-47735215* for *Pgc* (mm10 coordinates) (Fig. 7d). Further analysis of *Arid5b* suggests that it functions as a novel regulator of chief cells. In silico perturbation of *Arid5b* caused cells to deviate from the chief state, exhibiting pronounced trajectories across multiple directions. This perturbation also resulted in the downregulation of *Bhlha15* and *Zbtb16*, key TFs within the chief eGRN, and concomitant upregulation of *Capn8*, a marker characteristic of foveolar cells (Fig. 7e).

Unlike highly specific eGRNs of foveolar, parietal and chief cells, the mucous neck cells were characterized by less pronounced transcriptional networks, despite strong expression of *Muc6,* a key marker of mucous neck cells (Fig. 1c). The putative regulatory network of neck cells was composed of *Etv4, Etv5, Sox5, Sox9 and Rarg* (Fig. 8d). *Rarg* has been linked with the expression of *Muc6* via the chr7:141634859-141635359 regulatory region, which was further confirmed after *Rarg* in silico perturbation led to a downregulation of *Muc6* across all cell types. Nonetheless, *Rarg* does not seem to regulate the expression of any TF involved in the neck eGRN, as its knockout did not affect their expression (Supplementary Fig. 3b).

**Figure 8.**
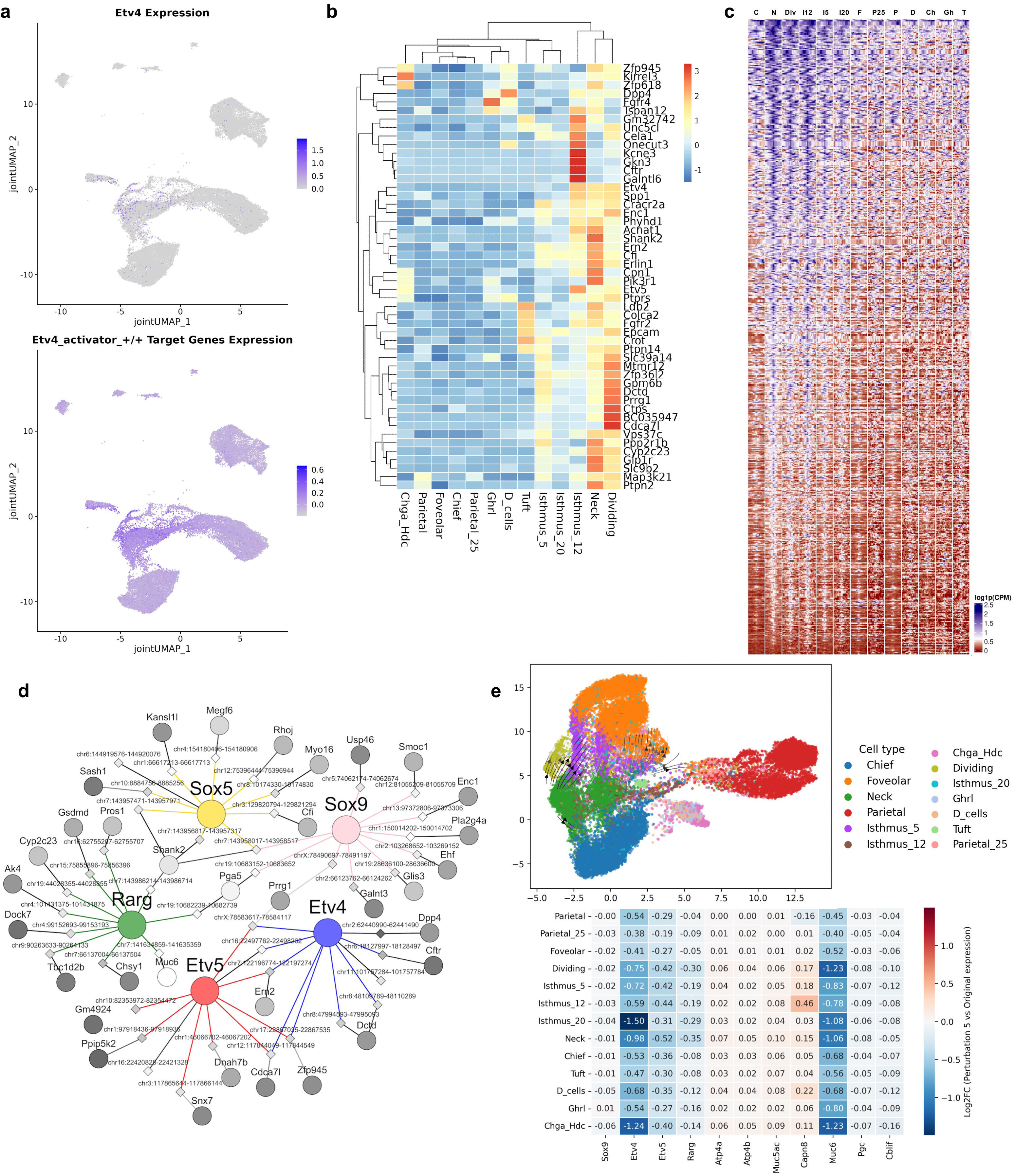
Neck cell regulatory program recovering the role of Etv4 and suggesting Sox5 as a novel regulator. **a.** UMAP displaying Etv4 expression and the expression of its target genes across epithelial clusters. **b.** Heatmap showing expression of top 50 target genes based on TF-to-gene importance values in the Etv4 direct activator eRegulon. **c.** Heatmap showing accessibility of all regulatory regions in the neck eGRN across cell types. **d.** Visualization of the eGRN of neck cells limited to the top 10 edges per TF based on triplet ranking values. Colorful circles represent TFs, diamonds represent enhancer regions and smaller circles represent target genes. Region-to-gene edges are shaded by their importance scores (importance_R2G) and nodes are shaded by log2FC values of expression and chromatin accessibility, where darker shades indicate higher values. **e.** Visualization of the effect of Sox5 in silico perturbation simulation after 5 iterations. Projected effect in the SCENIC+ UMAP embedding where arrows indicate each cell’s shift in embedding space, shaded by distance traveled, with darker arrows marking larger movements. Heatmap displaying log2FC between original gene expression and perturbed expression across epithelial cell types and genes of interest.

In silico perturbation demonstrates a key role of *Sox9* consistent with literature identifying it as an essential master regulator of neck cells, driving their differentiation and cell identity^29^. In silico knockdown of *Sox9* triggered downregulation of *Muc6, Etv4, Etv5* and *Rarg* expression, placing it at the center of this regulatory network. It further triggered putative induction of isthmus and chief cells states (Supplementary Fig. 3c). Similarly to the foveolar cell types, the eGRNs describing neck cells were already present in the isthmus and dividing cell populations. This was demonstrated by the expression patterns of the top 50 target genes within the *Etv4* eRegulon (Fig. 8b), and it was especially apparent in the chromatin states (Fig. 8c). On the other hand, the role of *Sox5* in the gastric epithelium during homeostasis is not very well understood. Compared to *Sox9* perturbation, in silico knockout of *Sox5* resulted in similar yet more pronounced gene expression changes, displaying a stronger downregulation of *Muc6, Etv4, Etv5* and *Rarg*. It also caused a slight upregulation of foveolar marker *Capn8* in isthmus_12 (Fig. 8e), which was not seen with *Sox9* perturbation (Supplementary Fig. 3c). Interestingly, perturbation of *Sox5* did not affect *Sox9* expression, and vice-versa, suggesting that both factors may cooperate in maintaining neck cell identity. Simultaneous perturbation of *Sox5* and *Sox9* showed similar but stronger effects (Supplementary Fig. 4b).

In addition to the aforementioned regulatory networks, we further observed a small eGRN with *Nfia* at its center that was linked to the *Nfib* and *Nfix* TFs. Unlike all other eGRNs, *Nfia* activity was linked with neck, chief and parietal cells. Its in silico knockdown suggested that its role is essential for the maintenance of neck, chief and parietal cells. Upon knockdown, the cellular states shifted away from these cell types and moved towards isthmus and foveolar cell types as supported by an upregulation of foveolar marker genes (*Muc5ac, Capn8*) in most cell types but with a more pronounced effect in isthmus_12 and neck cells (Supplementary Fig. 3d). Combined perturbation of *Nfia, Nfib* and *Nfix* amplified the effects observed in the *Nfia-*only knockout (Supplementary Fig. 4c).

### Isthmus cells are highly plastic cell types with stem-like properties

Isthmus clusters defined in this study (Isthmus_5, 12 and 20) sit between the foveolar and neck clusters in the WNN UMAP (Fig. 1b and 2a) and in the SCENIC+ UMAP (Fig. 2b), following previously published data of progenitor cells in the mouse stomach^40^. Isthmus clusters also express gene markers of foveolar (*Muc5ac*), neck (*Muc6*), and chief cells (*Pgc*) (Fig. 1c). Further investigation of the enrichment of eGRNs in the isthmus cells did not yield conclusive regulatory networks. We observed that all isthmus subclusters are defined by a regulatory overlap of eGRNs strongly enriched in foveolar, neck and chief cell types. This was particularly pronounced within the chromatin states. For example, the isthmus cell types had open chromatin at the regulatory regions defining *Sox5, Sox9* (neck), *Nfia* (chief and parietal) and *Klf4* (foveolar) (Fig. 3b).

MultiVelo analysis uncovered two differentiation patterns beginning at various isthmus cell clusters and progressing towards chief cells via neck cells or toward foveolar phenotypes (Fig. 2c,d). Similar to the MultiVelo results, cellular plasticity scores showed that isthmus cell types are the most plastic, with fixation of cellular states as they differentiate towards known mature cell types such as parietal, chief and foveolar (Fig. 2e). We also observed the highest EpiCHAOS score in the isthmus and foveolar cell types (Fig. 2f). We reasoned that the diversity of epigenetic states might be a feature of stem-like or rapidly changing cellular states. Taken together, our analysis suggests that isthmus cell types are transcriptionally and epigenetically the most diverse and “disorganized” (or plastic) of all stomach epithelial populations. They are associated with cellular division suggesting stem-like properties on the genome and transcriptome-wide scale.

Next, we investigated the relationship between the dividing cell types (*Mki67*) with other epithelial cell types. UMAP projection demonstrated that the dividing cells (Supplementary Fig. 5c) comprise their own cell cluster on the expression-based projection but are intermixed with the isthmus cell types (Supplementary Fig. 5d,e,f) on the chromatin state-based projection, further supporting the stem-like properties of isthmus cell types. Dividing cells also show the highest plasticity score after isthmus clusters, confirming the idea that these cells originate from the isthmus compartment in the healthy stomach (Fig. 2e).

GATA binding protein 4 (*Gata4*) is a pivotal regulator in mouse embryonic development and is the main regulator of columnar epithelium morphogenesis in the embryonic stomach^44,45^. *Gata4* direct activator eRegulon activity covered parietal_25, foveolar, dividing, isthmus_5 and isthmus_20 clusters as shown by target regions and target genes enrichment values (Fig. 3b). In silico knockout of *Gata4* led to an upregulation of stemness marker *Lgr5* across these clusters, as well as a downregulation of foveolar markers (*Capn8, Muc5ac*) and an upregulation of chief marker *Cblif* across all cell types (Supplementary Fig. 3e).

### Specific programs governing tuft and enteroendocrine cells

Tuft cells are known to depend on the TF *Pou2f3* for their differentiation and lineage specification^46^. Consistent with this, our analysis identified the extended activator *Pou2f3* eRegulon that was specific to tuft cells (Fig. 3a), supported by increased accessibility of its target regions and concomitant upregulation of its target genes (Supplementary Fig. 2b). Moreover, the *Ehf* direct activator eRegulon showed high enrichment values for tuft cells (Fig. 3b). Detailed examination of the target regions for the direct activator *Ehf* eRegulon and the extended activator *Pou2f3* eRegulon showed that both sets of regions are markedly more accessible in tuft cells than in the other epithelial populations (Supplementary Fig. 6a).

A subset of direct activator eRegulons was also associated with enteroendocrine cell populations, such as *Klf12, Rfx3* and *Rfx6* for Ghrl cells and *Bach2, Ebf1, Ets1, Lhx5, Rbpjl, Zfp641* for Chga_Hdc cells (Fig. 3a,b). Inspection of the target regions for eRegulons associated with the Ghrl (Supplementary Fig. 6b) and Chga_Hdc clusters (Supplementary Fig. 6c) showed that these loci are broadly accessible across enteroendocrine cells, with partial overlap in accessibility with the D_cells cluster.

## 3. Discussion

This study used scMultiome sequencing to identify the mechanisms underlying cell identity in the mouse stomach during homeostasis by constructing transcriptomic and epigenomic profiles of all cell types. This technology makes it possible to directly associate regulatory regions with their downstream gene activity, offering a clearer view of the molecular programs shaping cellular behavior^47^. The joint profiling of gene expression and chromatin accessibility is key to better characterize stem-like cell types that feature distinct transcriptomic and epigenomic profiles that change dynamically with their differentiation potential.

We extended our analysis by constructing cell-type specific eGRNs, identifying potential TFs and candidate enhancers that act as regulators and enhancer regions for gastric epithelial cell types. We used the inferred networks as a framework to perform in silico perturbation experiments where the expression of TFs of interest was set to 0 and the perturbed gene expression values propagated over five cycles to model indirect regulatory shifts occurring through hierarchical TF-to-TF interactions^39^. Multiple tools were further applied to characterize dynamic cell-state behavior. MultiVelo^36^ was used to predict cell fate trajectories through the assessment of latent time, EpiCHAOS^38^ was used to measure epigenetic diversity within cell types and epigenetic plasticity score^37^ was used to estimate how easily a cell might shift phenotypes by assessing the uncertainty in linking its chromatin landscape to its gene-expression profile.

Our analysis recovered three different isthmus populations (Isthmus_5, Isthmus_12 and Isthmus_20) that display many overlapping genes expressed at different proportions, indicating potential subpopulations of isthmus stem cells in the healthy stomach. Isthmus clusters are highly plastic, displaying non-specific and heterogeneous chromatin accessibility patterns that overlap with patterns displayed by foveolar, neck and chief cells. This indicates that during homeostasis, isthmus progenitor cells in the mouse stomach neither feature specific regulators, nor show a specific pattern of gene expression or chromatin accessibility. These cells, on the other hand, are highly heterogeneous and primed at the chromatin level to become different epithelial cell types depending on fate decisions.

We identified a new cell population, parietal_25, that shows transcriptomic profiles like parietal cells, but their epigenetic profile is similar not only to parietal but also foveolar and isthmus cells. This suggests that this population originates from isthmus cells and upon early expression of cell-type specific regulators, it commits to either a parietal or foveolar cell fate.

A recent study uncovered the role of TF *Esrrg* in the differentiation of stem cells into parietal cells^27^. Our study confirmed this role and further characterized the cooperativity between *Esrra, Esrrb* and *Esrrg* in the eGRN of parietal cells. We also identified *Ppara* as a new regulator of parietal cell identity. It is expressed in many tissues such as the liver, kidney, heart and intestine and plays a role in cellular processes such as proliferation, differentiation and apoptosis^42^. Nevertheless, the role of *Ppara* in the stomach across various species is limited, with no studies related to parietal cells in the existing literature.

Similarly, our in silico perturbation analyses corroborated the role of *Klf4* in maintaining foveolar cell type identity^28^. We further identified *Pparg* as a new putative regulator of foveolar cells, displaying an extensive effect on all TFs involved in the foveolar eGRN and on foveolar marker genes, where its knockout caused foveolar cells to shift states and lose their foveolar identity.

*Bhlha15* emerged as one the main TFs in the chief eGRN, in agreement with the literature^16^. Our in silico knockdown analyses further validated it as a key determinant of chief identity, together with a specific enhancer region linked to the expression of *Cblif*, a marker of this lineage. We also identified *Arid5b* as a previously unrecognized TF functioning as a regulator of chief cells, a finding that was supported by our in silico perturbation analyses.

*Sox5*, along with *Sox9,* was identified as a regulator of neck cells in the current study. The pathological role of *Sox5* upregulation has been extensively studied and it is shown to promote migration, invasion and metastasis in gastric cancer cells and other cancer types^48,49^. However, there is currently a lack of understanding of its role in gastric epithelial cell differentiation under homeostatic conditions. We show cooperation between *Sox5* and *Sox9* in the healthy stomach and demonstrate a synergistic role in maintaining neck cell identity.

Conceptually, our data support the notion of “substates and attractors” as a model describing the relationship between the stem cells and differentiated cell types^50^. Our analysis, for the first time outside of hematopoiesis, indicates that differentiated cell types can be described by clear, highly expressed cell states defined by cell type specific eGRNs (“attractors”). Conversely, the stem-like isthmus cell populations exist in a fluid, highly plastic state composed of many “substates” that can, but do not have to, drive stem cells towards “attractor” states.

Crucial to this conclusion is our observation that isthmus cells contain epigenetic states of the eRegulons of almost all differentiated cell types, even though they are not exhibiting the transcriptional profiles of these eRegulons. The epigenetic plasticity of isthmus cells primes them towards all fates that are upon activation of specific eGRNs, committing them towards specific, differentiated cell types. Future work is required to delineate the factors necessary to trigger cellular state commitment.

A limitation of this work is the use of computational models to infer enhancer-gene regulatory networks, latent temporal trajectories, plasticity scores and perturbation responses. While these tools offer significant insights into the regulatory logic of cellular states, our findings represent putative regulatory relationships rather than direct evidence of causality. Furthermore, the identity and lineage potential of parietal_25 and isthmus populations require further experimental validation.

In conclusion, this study uncovered previously unrecognized regulatory features, including potential regulators and cell type-specific programs, thereby providing new insight into the transcriptional and epigenomic organization of the healthy gastric epithelium. Despite these advances, experimental validation of the identified TFs in healthy mouse gastric tissue remains necessary to confirm the functions of these regulators and their associated enhancers, and to clarify their contributions to cell-fate specification and cellular identity.

## 4. Methods

### 4.1 Sample preparation

Murine stomach tissue was collected from healthy wild-type C57BL/6 mice (N=10, 5M, 5F) of 8-12 weeks of age. Animals were euthanized in accordance with Columbia University’s approved Institutional Animal Care and Use Committee (IACUC) protocol (AC-AACB3701) using an approved euthanasia method. All procedures were conducted in compliance with applicable animal welfare regulations. Immediately following euthanasia, the abdominal cavity was opened under sterile conditions, and the stomach was collected and then rinsed in ice-cold sterile PBS to remove luminal contents and blood. Excess connective tissue was trimmed, and the stomach was cut along the greater curvature and placed on ice prior to freezing.

Specimen storage containers were pre-cooled by exposure to liquid nitrogen (LN₂) vapor prior to tissue freezing. To generate a stable LN₂ vapor phase, a benchtop Dewar flask (Thermo Fisher Scientific, Cat. No. 4150-1000) was filled with liquid nitrogen, and a 250 mL stainless steel beaker (Ace Glass Incorporated; VWR, Cat. No. 8671141) was suspended within the Dewar such that it was surrounded by LN₂ vapor but not in direct contact with the liquid phase. A layer of aluminum foil was placed at the bottom of the stainless-steel beaker to serve as a thermal interface. Dissected stomach tissue samples were placed directly onto the foil surface at the bottom of the stainless steel beaker inside the Dewar specimen storage container and exposed for 2 minutes to LN₂ vapor for rapid freezing.

Following complete freezing, samples were removed from the beaker, tightly wrapped in aluminum foil, sealed in biohazard ziplock bags, and stored at −80 °C. Frozen samples were transported to the New York Genome Center on dry ice under continuous cold-chain conditions to preserve sample integrity.

### 4.2 Multiome library preparation and sequencing

#### Mouse Tissue Nuclei Isolation and Hashing

Frozen mouse gastric tissue weighing 50 mg was quickly minced on ice with 200 μL of chilled Homogenization Suspension Buffer (HSB: 320 mM Sucrose, 10 mM Tris-HCl pH 7.4, 3 mM CaCl_2_, 3 mM Mg(Ac)_2_, 1 mM DTT and 1 U/µL RNase Inhibitor) and transferred to a low-bind microcentrifuge tube embedded in ice. 0.15% NP40 was added to the minced tissue and a 5 minute lysis was initiated. Tissue was dissociated with 10x-15x strokes of a disposable polypropylene pestle. An additional 300 μL of HSB and 0.15% NP40 was added to the tissue to obtain a total volume of 500 μL, followed by pipet mixing 10x. The lysis reaction was quenched with 1 mL of HSB and the homogenate was filtered through a 30 μm filter. An additional 4 mL of HSB was added to wash the filter and optimize nuclei recovery.

The nuclei filtrate was centrifuged at 500g for 5 minutes at 4°C and the nuclei pellet was resuspended in 200 μL of Nuclei Staining buffer (NSB: Cell Staining Buffer from BioLegend, 0.9 mM CaCl_2_, 0.5 mM MgCl_2_, 0.01% Tween-20, 1 mM DTT and 1 U/µL RNase Inhibitor). Nuclei quantity was assessed using trypan blue stain and a Countess II Automated Cell Counter. For the sample hashing step, 1 million nuclei were transferred to a new 1.5 mL low-bind microcentrifuge tube and the suspension volume was brought up to 100μL with NSB. The nuclei suspension was blocked on ice with Mouse TruStain FcX™ blocking solution (BioLegend) for 10 minutes, after which a unique hashtag oligonucleotide antibody (HTO: TotalSeq™-A anti-Nuclear Pore Complex Proteins Hashtag Antibodies from BioLegend) was added to achieve a hashing concentration of one million nuclei/1 µg HTO/100 µL.

Hashing incubation proceeded for 30 minutes, followed by 3 washing steps with NSB. Nuclei quality and clumping conditions were quickly assessed under a fluorescent microscope (Nikon ECLIPSE Ti2-E) and a 20 μm filtration was performed if clumps greater than 30 μm constituted more than 10% of the nuclei suspension. Nuclei quantity was assessed as described above.

#### Nuclei Pooling for Multiome Assay and Sequencing

To proceed with the permeabilization step, samples with unique HTOs and at equal proportions were pooled together to a maximum of two million nuclei (Pool Batch 1: 6 samples, Pool Batch 2: 4 samples). The samples pool was mixed well, centrifuged at 500g for five minutes at 4°C, and the nuclei pellet was resuspended in Permeabilization Wash Buffer (PWB: 10 mM Tris-HCl pH 7.4, 10 mM NaCl, 3 mM MgCl_2_, 0.01% Tween-20, 1% BSA, 1 mM DTT and 1 U/µL RNase Inhibitor) for buffer exchange, followed by centrifugation at 500g for five minutes at 4°C. The supernatant was completely removed and permeabilization was initiated as soon as the nuclei pellet was resuspended with 125 µL of Permeabilization Buffer (PB: PWB, 0.015% NP-40, 0.015% Digitonin, 1 mM DTT and 1 U/µL RNase Inhibitor).

After the 2 minutes of permeabilization incubation, 1 mL of PWB was added to quench the reaction and the nuclei suspension was centrifuged at 500g for 5 minutes at 4°C. The permeabilized nuclei pool was resuspended in 125 µL of Nuclei Buffer (NB: 1× 10X Genomics Nuclei Buffer, Nuclease-free Water, 1 mM DTT and 1U/µL RNase Inhibitor), followed by nuclei quantification. To target a recovery of 2500-5000 nuclei per sample (sample condition dependent), 4-4.25×10^4^ nuclei (8-8.5 million nuclei/mL) were used to proceed with the transposition step as per 10X Genomics Multiome protocol (Chromium Next GEM Single Cell Multiome ATAC + Gene Expression Reagent Kits User Guide, CG000338 Rev G, 10X Genomics). To generate GEX, ATAC and HTO libraries, 25 µL of the pre-amplified product was used as input for each library.

The resulting libraries were sequenced on a NovaSeq 6000™ (Batch 1) and NovaSeq™ X Series 25B (Batch 2) flow cell (Illumina), with a target of 25000 reads/cell for GEX and ATAC libraries and 3000 reads/cell for HTO library. GEX and HTO libraries were sequenced together in the same lane (sequencing parameters: 28 bp read 1, 10 bp i7 index, 10 bp i5 index, 91 bp read 2), whereas ATAC libraries were sequenced separately in their own lane (sequencing parameters: 50 bp read 1, 8 bp i7 index, 24 bp i5 index, 50 bp read 2).

### 4.3 Multiome data processing and analyses with Seurat/Signac

Cell Ranger ARC was used to process multiome chromium single nuclei ATAC and gene expression sequencing data (cellranger-arc-mm10-2020-A) with manual editing to bypass the 20,000 nuclei limit (“MAX_CELLS = 20000”). Reads were then mapped to the mouse reference genome (GRCm38 - 2020-A). Cell Ranger ARC outputs were imported to R using Seurat^30^ (version 5.3.0) and Signac^31^ (version 1.14.0). MACS3^51^ was used within Signac to recall peaks while removing peaks mapped to genomic blacklist regions from the mouse genome. Datasets were filtered based on the snRNA-seq quality where cells that expressed fewer than 100 genes and over 10% counts aligning to mitochondrial genes were removed. Cells with fewer than 1,000 and over 100,000 fragments, transcription start site < 1, percentage of reads in peaks < 15% were also removed based on the snATAC-seq assay.

HTO libraries were processed separately using the Kallisto/Kite workflow to generate hashtag counts and perform sample demultiplexing^52^. Singlets were kept using a modified version of Seurat *HTODemux* with a hashtag library-size normalization modification to increase sensitivity. After quality control, a total of 31,598 cells were analyzed.

The snRNA-seq data for each sequencing dataset was individually pre-processed using Seurat by normalizing cells using *SCTransform* followed by principal component analysis (PCA) with the *RunPCA* function. The first 50 principal components (PCs) were used for dimensionality reduction and the Louvain algorithm to cluster cells with the *RunUMAP, FindNeighbors* and *FindClusters* functions. Similarly, the snATAC-seq data for each sequencing dataset was processed individually using Signac. Peaks were normalized using term frequency inverse document frequency (TFIDF) with the *RunTFIDF* function, peaks for dimensional reduction selected with *FindTopFeatures* and dimension reduction done using a singular value decomposition (SVD) using the *RunSVD* function. Graph-based clustering was performed using 50 latent semantic indexing (LSI) followed by FindNeighbors and FindClusters functions using the four resolutions previously described. WNN using the FindMultiModalNeighbors() function allowed for joint clustering of the RNA and ATAC assays for each sequencing batch separately, resulting in two pre-processed multiome objects. WNN represents a weighted combination of both RNA and ATAC modalities.

Finally, the RNA assays from both pre-processed multiome objects were merged with the *merge* function and processed as previously described (*SCTransform, RunPCA, FindNeighbors, FindClusters, RunUMAP*) and integrated with Harmony with the *IntegrateLayers* function (method = HarmonyIntegration).

A unified set of peaks was created and quantified across both ATAC assays using the *reduce* function from the GenomicRanges package and quantified with the function *FeatureMatrix*. The objects, containing ATAC assays with the same features, were then merged with the *merge* function and processed as previously described (*FindTopFeatures, RunTFIDF, RunSVD, FindNeighbors, FindClusters, RunUMAP*). Both ATAC assays were then integrated with the *IntegrateEmbeddings* function, using common features across selected with the *FindIntegrationAnchors* function. The integrated assays were processed as previously described and then integrated together using WNN with the *FindMultiModalNeighbors* function, followed by *RunUMAP* and *FindClusters* at four different resolutions (0.2, 0.5, 0.8, 1.0).

Differential gene expression analysis was performed by creating an aggregate expression per animal per cluster using the *AggregateExpression* function (assay = “SCT”, slot = “counts”, group.by = c(“HTO_origin”, “wsnn_res.1”)), followed by *FindAllMarkers* with DESeq2 (test.use = “DESeq2”) to explore differentially expressed genes (DEGs) across clusters. Cell annotation was performed manually based on the expression of known markers and based on the top DEG per cluster. Epithelial cells (Parietal, Parietal_25, Foveolar, Dividing, Isthmus_5, Isthmus_12, Isthmus_20, Neck, Chief, Tuft, D_cells, Ghrl and Chga_Hdc clusters) were subsetted resulting in 27,810 cells, 514,311 regions and 24,596 genes that were used as an input for SCENIC+ to uncover gene regulatory networks.

### 4.4 Gene regulatory network analysis and in silico perturbation with SCENIC+

The processed snRNA-seq Seurat object with cell type annotation was converted to a h5ad file and used as input for the SCENIC+ analysis. For the snATAC-seq data, the raw fragment files were used to preprocess the data using pycisTopic as described by the authors^39^. During snATAC-seq data preprocessing, topic modelling was performed with MALLET where models with 30, 35, 40, 45, 50, 60, 70, 80, 90, 100 and 110 topics were tested. Based on the model selection metrics, the model with 80 topics was selected.

Prior to running SCENIC+, a custom cistarget database was created using the *create_cisTarget_databases* Python package using the ATAC-seq consensus peaks calculated per cell type and a previously annotated collection of motifs made available by the SCENIC+ authors as input. SCENIC+ was run using default parameters targeting 514,311 regions and 24,596 genes in a total of 27,810 cells. A total of 468 eRegulons were produced by the SCENIC+ snakemake pipeline, where 170 were direct and 298 were extended. The AUCell algorithm^53^ is used within the pipeline to calculate AUC (area under the curve) scores of target genes and target regions for each eRegulon.

Genes in the dataset were considered TFs if included in a list made available by the authors on https://resources.aertslab.org/cistarget/tf_lists/allTFs_mm.txt, also available in Supplementary Table 2. Direct eRegulons are inferred on annotation of TFs based on experimental mouse ChIP-seq data and extended eRegulons are inferred through human orthologs or motif similarity^39^.

A downstream filtering step was applied to the eRegulons using the “*apply_std_filtering_to_eRegulons*” function as a quality control to remove negative region-to-gene correlations (repressive regulatory relationships between regions and genes) and only keep extended eRegulons if no direct version exists to prevent redundancy. Annotation “+/+” of eRegulons represent TFs that activate gene expression while opening chromatin, whereas “-/+” eRegulons represent TFs that repress gene expression despite having open chromatin regions at their target genes.

AUC values represent the enrichment score of eRegulons in individual cells, calculated using the AUCell algorithm. Regulon specificity score (RSS) values measure how specifically an eRegulon is active in particular cell types and are calculated using Jensen-Shannon divergence. AUC and RSS values were used to link eRegulons to cell types and only direct activator (“+/+”) eRegulons were used in the construction of cell type-specific eGRNs.

In silico perturbation analyses were performed using the predictive modeling framework implemented in SCENIC+ using default parameters, which leveraged the inferred eGRN as the basis for feature selection and regulatory inference. For each gene in the eGRN, a regression model was trained to predict expression from its upstream TFs, restricted to those with enhancer-linked binding sites. TF knockdowns were simulated by setting the expression of the targeted TFs to zero and iteratively propagating predicted expression values through the network for five cycles to capture indirect regulatory effects. The resulting simulated expression matrix was co-embedded with the original UMAP to visualize predicted cell-state shifts. A delta-embedding quantified the direction and magnitude of the perturbation-induced transitions, enabling identification of TFs driving specific lineage changes.

### 4.5 Trajectory analysis with MultiVelo

MultiVelo was used to infer latent time and identify early cells along the differentiation trajectory of gastric epithelial cells^36^. To obtain spliced and unspliced RNA counts, velocyto^54^ was run on the original BAM files using default parameters. To integrate chromatin accessibility with transcriptional dynamics, a linked peaks matrix was generated from the ATAC-seq and RNA modalities using the *LinkPeaks* function in Signac with default settings. The resulting RNA velocity and linked peaks outputs were imported into Python as AnnData objects and pre-processed following the recommendations in the MultiVelo documentation.

Briefly, the RNA modality was filtered, normalized, and highly variable genes were selected using scVelo (*scv.pp.filter_and_normalize*) with min_shared_counts = 10 and n_top_genes = 3000. No additional preprocessing was applied to the ATAC modality, as this had already been performed in Seurat. Both modalities were subsequently subset to retain overlapping cells and features. RNA neighborhood connectivities were computed using scVelo (*scv.pp.moments*) with n_neighbors = 50. WNN graphs were recalculated in Seurat, as recommended by the MultiVelo authors. MultiVelo was then run using the dynamic model via *mv.recover_dynamics_chrom* with default parameters. Latent time was computed using *mv.latent_time*.

All analyses were performed separately for each batch. Per-cell latent time estimates derived from Batch 1 and Batch 2 were added to the integrated Seurat object via cell barcode matching. Latent time values were subsequently visualized on the WNN UMAP embedding to enable comparison of differentiation progression across datasets.

### 4.6 Epigenetic plasticity scores

Epigenetic plasticity was computed as previously described^37^. Briefly, the RNA modality was normalized and log-transformed using Scanpy (*sc.pp.normalize_total, sc.pp.log1p*), and highly variable genes were selected with *sc.pp.highly_variable_genes* (n_top_genes = 3000, flavor = “seurat”, subset = True). PCA was subsequently performed using sc.pp.pca.

For the ATAC modality, a gene activity score matrix was computed in Signac using the *GeneActivity* function with default parameters. The resulting matrix was imported into Python as an AnnData object, normalized and subjected to PCA using the same Scanpy workflow as applied to the RNA modality. Both modalities were then restricted to overlapping features. To generate metacells, cells were grouped by cell type, batch, and sample. Groups containing more than 40 cells were further subdivided using Leiden clustering (*sc.tl.leiden*) with a resolution of 0.5. This procedure yielded a total of 492 metacells, representing approximately 1.8% of the original number of cells.

Average gene expression (RNA) and average gene activity scores (ATAC) were calculated for each metacell. Epigenetic plasticity was quantified by fitting a logistic regression model with a multinomial likelihood to predict cell type identity based on RNA gene expression. Shannon entropy was then computed across the predicted cell-type probability distributions for each metacell. Higher entropy values indicate greater disagreement between transcriptional and chromatin accessibility profiles and thus increased epigenetic plasticity. Finally, metacell-level plasticity scores were assigned back to individual cells belonging to each metacell.

## 4.7 Data availability

Raw (FASTQ) and processed (Cell Ranger) sequencing data will be deposited in Gene Expression Omnibus (GEO) repository. All code used for the analyses is available on GitHub (https://github.com/GastroEsoLab/scMultiome_Mouse_Gastric_Manuscript/).

## 5 Acknowledgements

We thank Will Liao and Ali Oku from the Computational Biology group at the New York Genome Center for the helpful discussions on the methodological aspects of SCENIC+ and perturbation analysis.

## Grant Support

This research was supported by grant R01CA2722891 from the National Cancer Institute (NCI) of the National Institutes of Health (NIH) to Sandra Ryeom, Silas Maniatis and Karol Nowicki-Osuch. This work was also made possible in part by the MacMillan Family Foundation as part of the MacMillan Center for the Study of the Non-Coding Cancer Genome at the New York Genome Center.

## Abbreviations

AUC: Area under the curve
DEGs: Differentially expressed genes
eGRN: Enhancer-driven gene regulatory network
GC: Gastric cancer
GEO: Gene Expression Omnibus
HSB: Homogenization suspension buffer
HTO: Hashtag oligonucleotide
IACUC: Institutional animal care and use committee
LN₂: Liquid nitrogen
LSI: Latent semantic indexing
NB: Nuclei buffer
NSB: Nuclei staining buffer
PB: Permeabilization buffer
PCA: Principal component analysis
PCs: Principal components
PWB: Permeabilization wash buffer
RSS: Regulon specificity score
scMultiome: Single-cell multiome
snATAC-seq: Single-nucleus ATAC sequencing
snRNA-seq: Single-nucleus RNA sequencing
SVD: Singular value decomposition
TF: Transcription factor
TFIDF: Term frequency–inverse document frequency
WNN: Weighted nearest neighbors

## Disclosures

The authors disclose no conflicts.

## Author Contributions

Maithê Rocha Monteiro de Barros, DVM, PhD (Conceptualization: Supporting; Data Curation: Lead; Formal Analysis: Lead; Software: Lead; Validation: Lead; Visualization: Lead; Writing - original draft: Lead)

Katharina Bosch, PhD (Formal Analysis: Supporting; Software: Supporting; Visualization: Supporting)

Salima Soualhi, PhD (Investigation: Equal; Resources: Supporting)

Shirin Issa Bhaloo, PhD (Investigation: Equal; Resources: Supporting)

Thomas Chu (Investigation: Equal; Resources: Supporting)

Tanya Hemrajani (Investigation: Equal; Resources: Supporting)

Jin Cho, PhD (Investigation: Equal, Resources: Supporting)

Kurtay Ozuner (Investigation: Equal; Resources: Supporting)

Rui Fu, PhD (Data Curation: Supporting; Software: Supporting)

Heather Geiger (Data Curation: Supporting; Software: Supporting)

Nicolas Robine, PhD (Data Curation: Supporting; Resources: Supporting; Software: Supporting; Supervision: Supporting)

Jade E.B. Carter, MSc (Investigation: Lead; Resources: Lead; Supervision: Supporting; Project Administration: Supporting)

Silas Maniatis, PhD (Conceptualization: Equal; Funding Acquisition: Lead; Methodology: Equal; Project Administration: Equal; Resources: Lead, Supervision: Supporting)

Sandra Ryeom, PhD (Conceptualization: Equal; Funding Acquisition: Lead; Methodology: Equal; Project Administration: Equal; Investigation: Lead; Resources: Lead; Supervision: Supporting)

Simon Tavaré, PhD (Conceptualization: Supporting; Funding Acquisition: Supporting; Resources: Lead; Supervision: Lead; Validation: Equal; Writing - review & editing: Lead)

Karol Nowicki-Osuch, PhD (Conceptualization: Equal; Funding Acquisition: Lead; Methodology: Equal; Project Administration: Equal; Supervision: Lead, Validation: Equal; Resources: Lead; Visualization: Supporting; Writing - review & editing: Lead)

All authors had access to the study data and had reviewed and approved the final manuscript.

## Notes

https://github.com/GastroEsoLab/scMultiome_Mouse_Gastric_Manuscript/

### Competing Interest Statement

The authors have declared no competing interest.

