## Supplementary Figures for "Single-nucleus multiome sequencing identifies candidate regulators of mouse gastric epithelial homeostasis"

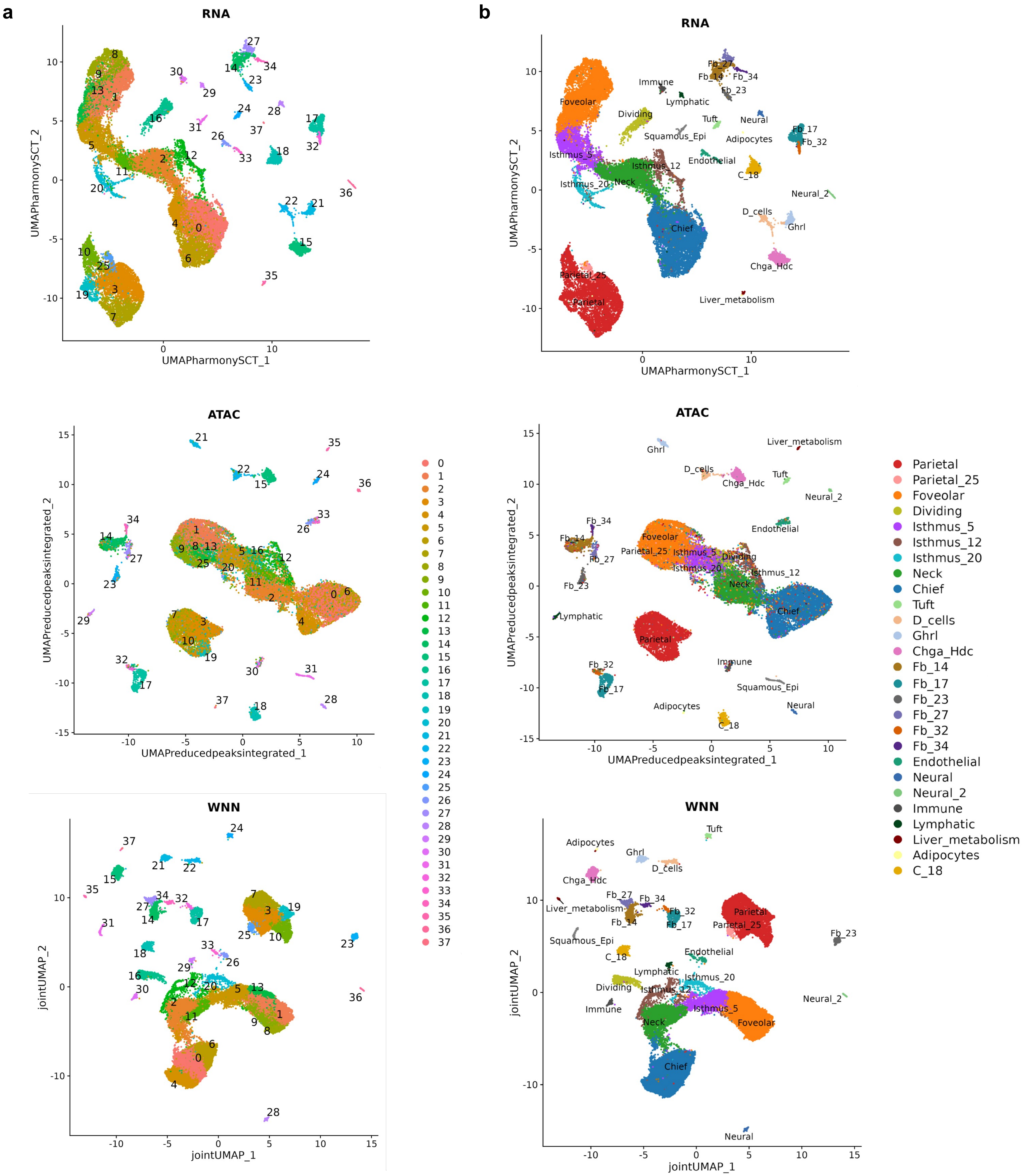


**Supplementary Figure 1 – Unsupervised and manually annotated cell clusters across RNA, ATAC and WNN embeddings.** *UMAP projections of all cells from RNA, ATAC, and WNN embeddings, shown with* ***a.*** *unsupervised cluster labels at resolution 1 (38 clusters) alongside* ***b.*** *manually curated cell-type annotations (27 cell types).*

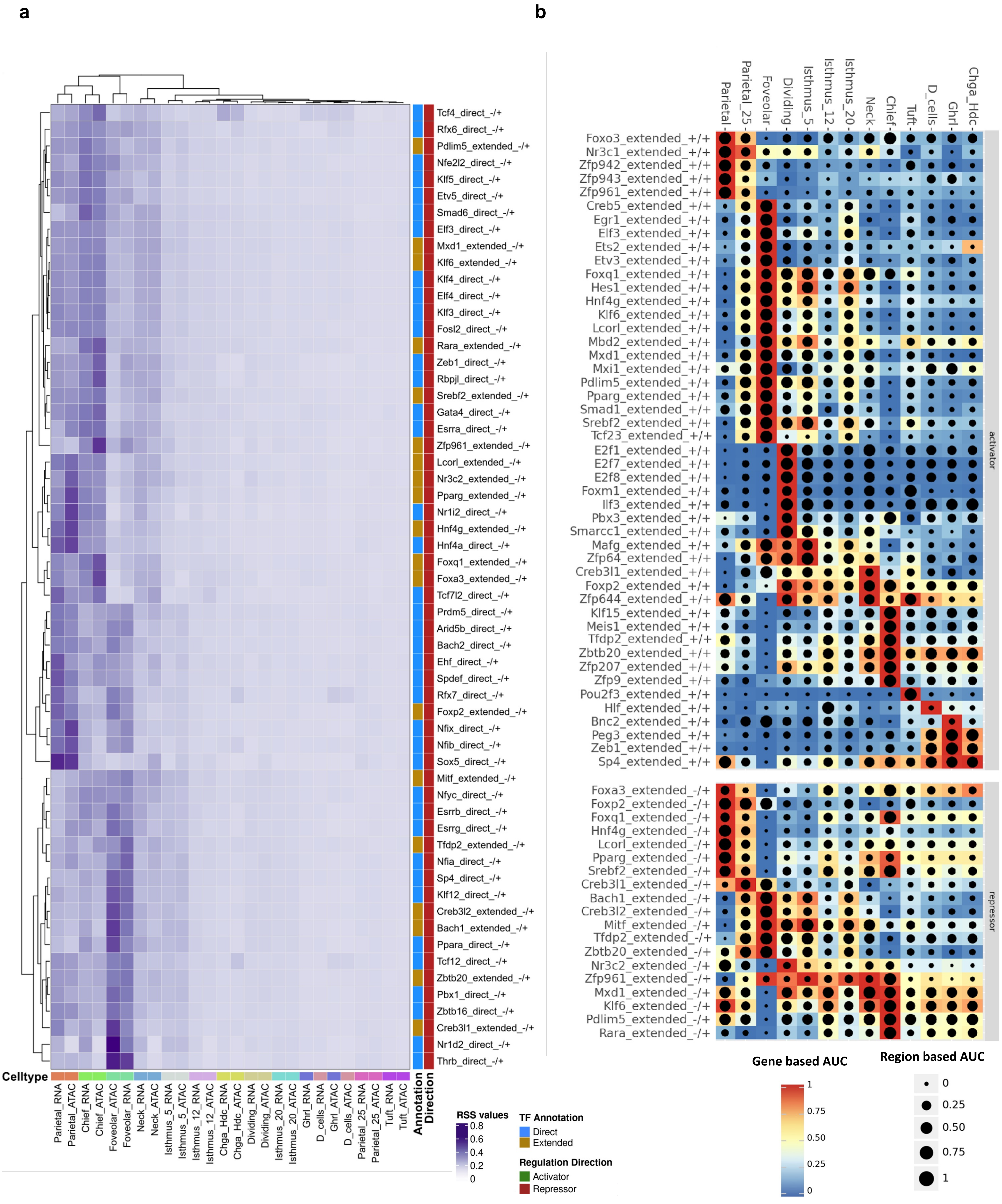


**Supplementary Figure 2 – SCENIC+ cell specificity scores (RSS) for all repressor eRegulons alongside eRegulon AUC enrichment scores for all extended eRegulons.** ***a.*** *Heatmap of Regulon cell specificity scores (RSS) for direct and extended repressor eRegulons across cell types based on the RNA and ATAC assays. Transcription factor (TF) annotation can be direct (blue) or extended (yellow). RSS values are calculated via Jensen-Shannon divergence to quantify the lineage-specific activity of each eRegulon. Heatmap is clustered by cell type (columns) and eRegulons (rows).* ***b.*** *Heatmap-dotplot of AUC gene (color) and AUC region (dot size) values per activator and repressor extended eRegulons across cell types. AUC values measure the enrichment of a regulon's target nodes within a cell's overall ranking of gene expression and chromatin accessibility. Cell types are ordered by transcriptomic similarity to highlight the dynamic shifts in regulatory networks.*

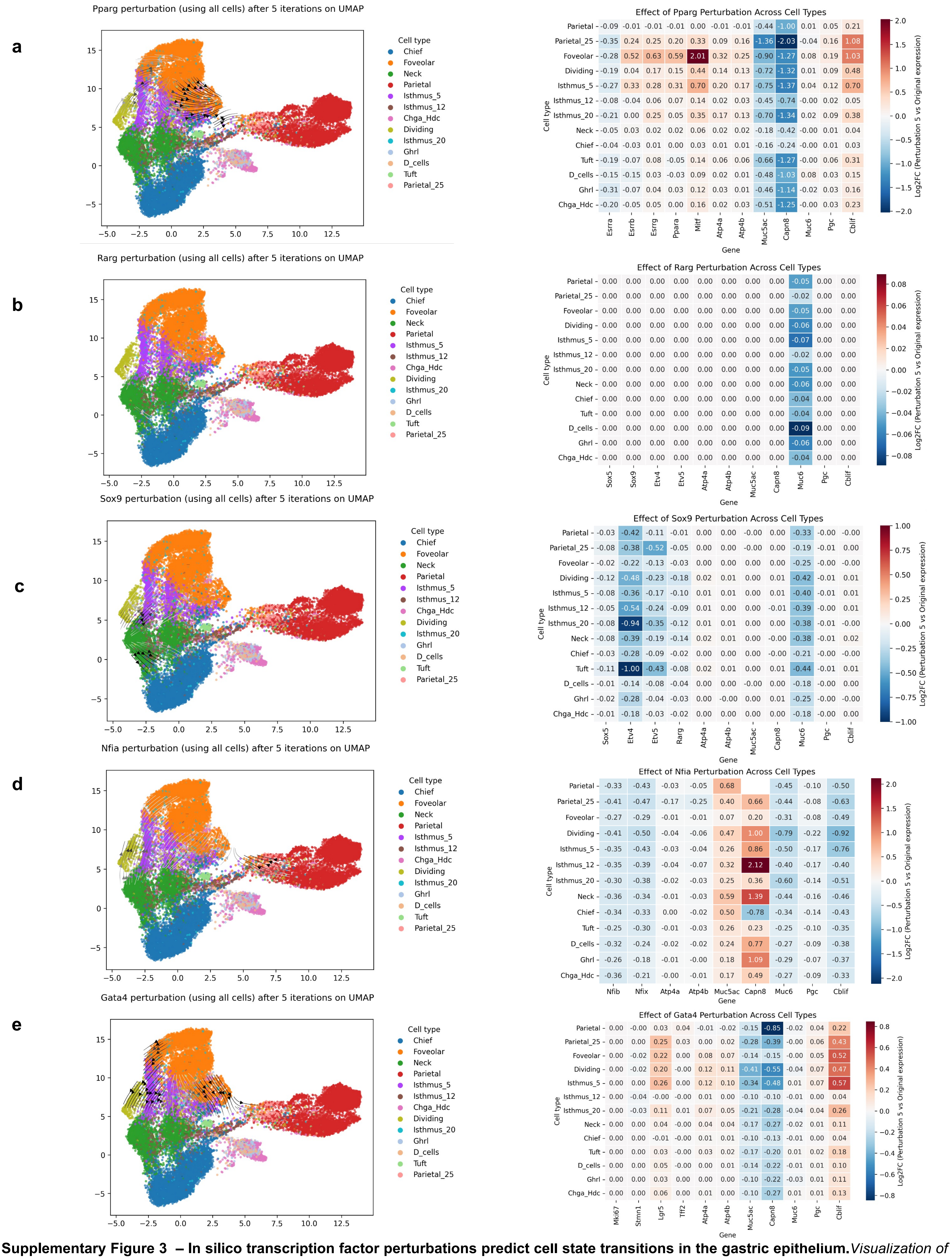


**Supplementary Figure 3 – In silico transcription factor perturbations predict cell state transitions in the gastric epithelium.** *Visualization of the effect of in silico perturbation simulation after 5 iterations for* ***a.*** *Pparg,* ***b.*** *Rarg,* ***c.*** *Sox9,* ***d.*** *Nfia,* ***e.*** *Gata4. Projected effect in the SCENIC+ UMAP embedding where arrows indicate each cell's shift in embedding space, shaded by distance traveled, with darker arrows marking larger movements. Heatmap displaying log2FC between original gene expression and perturbed expression across epithelial cell types and genes of interest.*

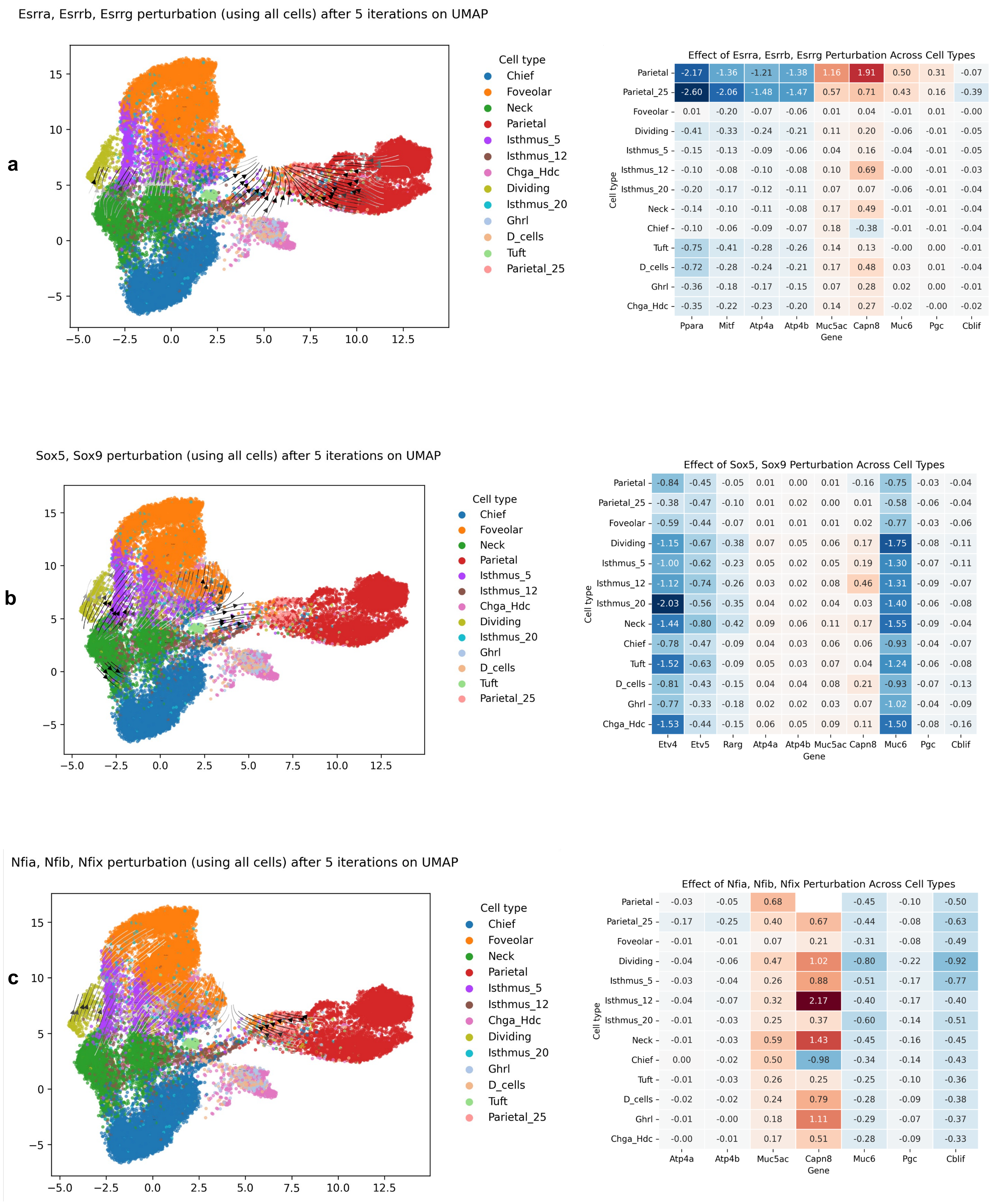


**Supplementary Figure 4 – Combined transcription factor knockdown simulations reveal cooperative regulation of gastric epithelial cell identity.** *Visualization of the effect of in silico perturbation simulation after 5 iterations for* ***a.*** *Esrra, Esrrb and Esrrg simultaneously,* ***b.*** *Sox5 and Sox9 simultaneously,* ***c.*** *Nfia, Nfib and Nfix simultaneously. Projected effect in the SCENIC+ UMAP embedding where arrows indicate each cell's shift in embedding space, shaded by distance traveled, with darker arrows marking larger movements. Heatmap displaying log2FC between original gene expression and perturbed expression across epithelial cell types and genes of interest.*

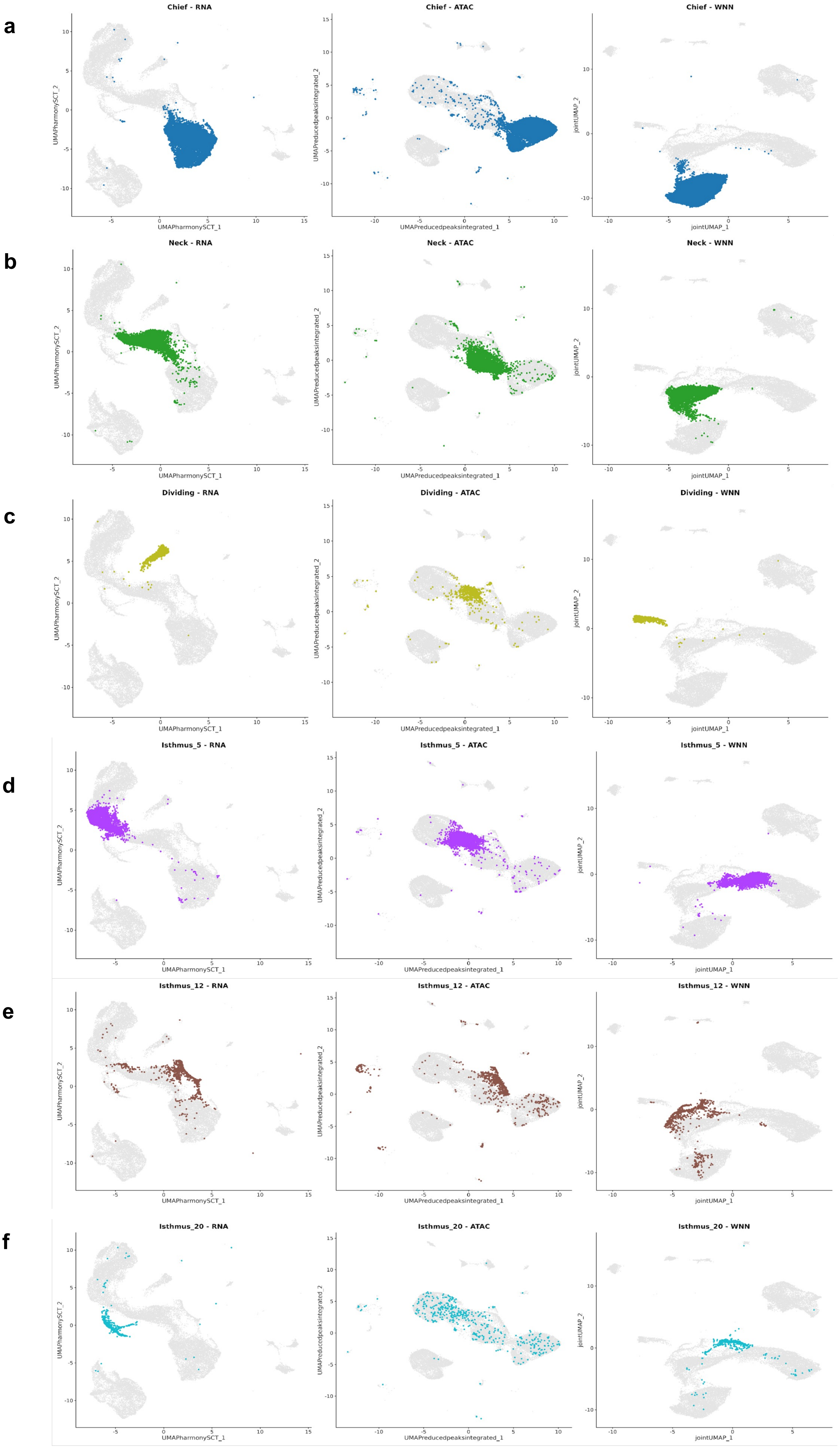


**Supplementary Figure 5 – Localization of gastric epithelial cell populations across multimodal embeddings.** *RNA, ATAC and WNN UMAP embeddings highlighting the distribution of* ***a.*** *Chief,* ***b.*** *Neck,* ***c.*** *Dividing,* ***d.*** *Isthmus_5,* ***e.*** *Isthmus_12 and* ***f.*** *Isthmus_20, demonstrating their transcriptional, chromatin-accessibility and integrated multimodal identities.*

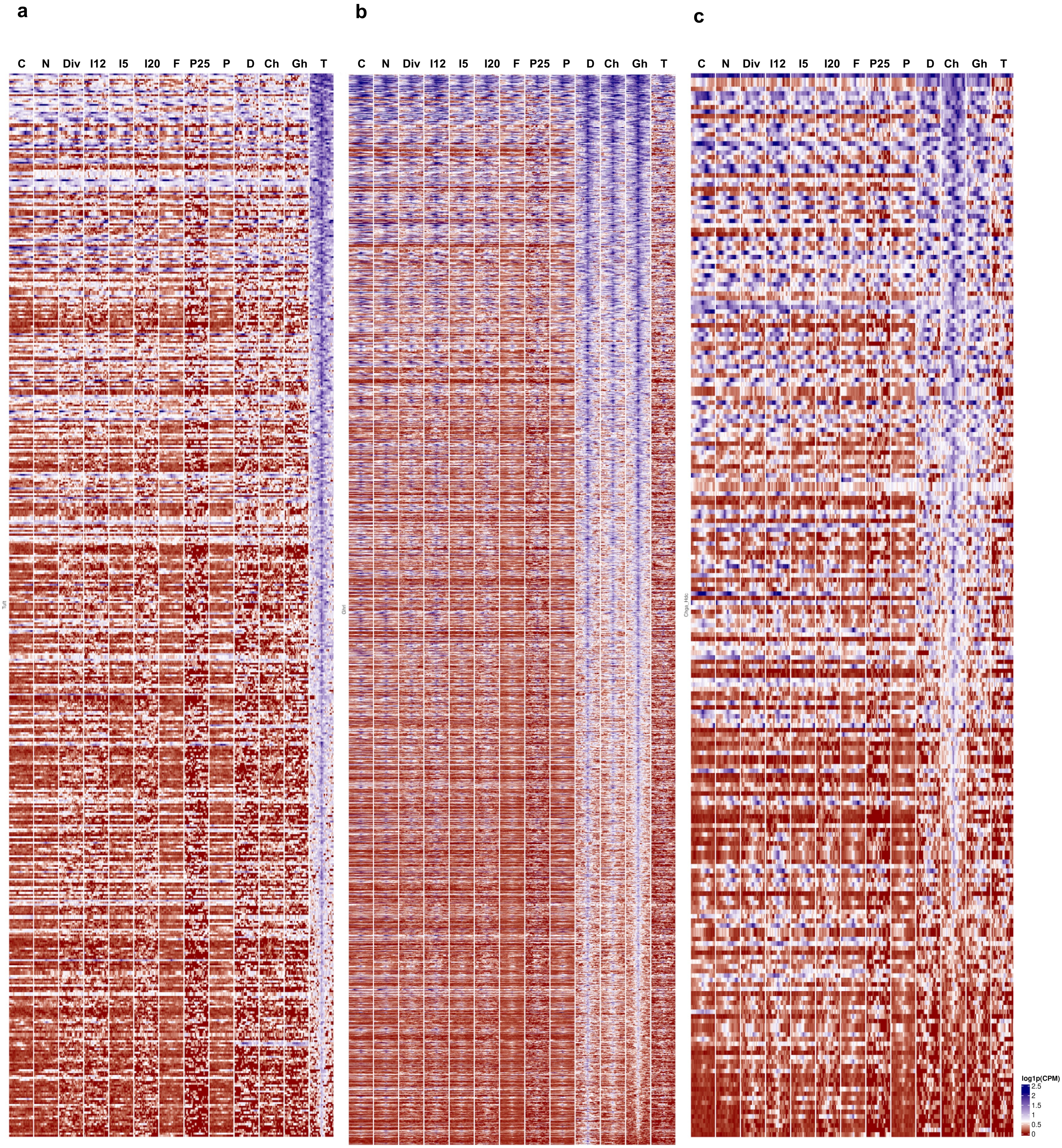


**Supplementary Figure 6 – Chromatin accessibility landscape of activator eRegulon target regions in tuft and enteroendocrine cells.** *Heatmap showing chromatin accessibility across all target regions assigned to direct activator eRegulons related to* ***a.*** *Tuft,* ***b.*** *Ghrl and* ***c.*** *Chga_Hdc cell types. C:Chief, N:Neck, Div:Dividing, I5:Isthmus_5, I12: Isthmus_12, I20:Isthmus_20, F: Foveolar, P25:Parietal_25, P:Parietal, D:D_cells, Ch:Chga_Hdc, Gh:Ghrl, T:Tuft cells.*
