## Supplementary Tables for "Single-nucleus multiome sequencing identifies candidate regulators of mouse gastric epithelial homeostasis"

**Supplementary Table 1 – Cellranger output information for Batch 1 and Batch 2 datasets**

| | **Batch 1** | **Batch 2** |
|---|---|---|
| Genome | mm10 | mm10 |
| Pipeline version | cellranger-arc-2.0.2 | cellranger-arc-2.0.2 |
| Estimated number of cells | 21860 | 22321 |
| Feature linkages detected | 1813529 | 1626190 |
| Linked genes | 18970 | 15599 |
| Linked peaks | 86019 | 103485 |
| ATAC Confidently mapped read pairs | 0.8675 | 0.9079 |
| ATAC Fraction of genome in peaks | 0.0291 | 0.0391 |
| ATAC Fraction of high-quality fragments in cells | 0.7642 | 0.8634 |
| ATAC Fraction of high-quality fragments overlapping TSS | 0.2732 | 0.3071 |
| ATAC Fraction of high-quality fragments overlapping peaks | 0.3702 | 0.4515 |
| ATAC Fraction of transposition events in peaks in cells | 0.322 | 0.4251 |
| ATAC Mean raw read pairs per cell | 134091.4 | 47533.15 |
| ATAC Median high-quality fragments per cell | 31355 | 15597 |
| ATAC Non-nuclear read pairs | 0.0103 | 0.0076 |
| ATAC Number of peaks | 95180 | 126119 |
| ATAC Percent duplicates | 0.5557 | 0.3747 |
| ATAC Q30 bases in barcode | 0.8876 | 0.8576 |
| ATAC Q30 bases in read 1 | 0.9463 | 0.9478 |
| ATAC Q30 bases in read 2 | 0.8793 | 0.9378 |
| ATAC Q30 bases in sample index i1 | 0.9446 | 0.859 |
| ATAC Sequenced read pairs | 2931238048 | 1060987358 |
| ATAC TSS enrichment score | 6.9178 | 9.912 |
| ATAC Unmapped read pairs | 0.0334 | 0.0099 |
| ATAC Valid barcodes | 0.9612 | 0.9547 |
| GEX Fraction of transcriptomic reads in cells | 0.8333 | 0.7326 |
| GEX Mean raw reads per cell | 118634.8 | 38989.27 |
| GEX Median UMI counts per cell | 7410 | 2511 |
| GEX Median genes per cell | 2869 | 1450 |
| GEX Percent duplicates | 0.8327 | 0.7818 |
| GEX Q30 bases in UMI | 0.9528 | 0.956 |
| GEX Q30 bases in barcode | 0.9552 | 0.9358 |

| | | |
|---|---|---|
| GEX Q30 bases in read 2 | 0.903 | 0.9359 |
| GEX Reads mapped antisense to gene | 0.0471 | 0.0441 |
| GEX Reads mapped confidently to exonic regions | 0.2969 | 0.3671 |
| GEX Reads mapped confidently to genome | 0.8543 | 0.847 |
| GEX Reads mapped confidently to intergenic regions | 0.1896 | 0.1465 |
| GEX Reads mapped confidently to intronic regions | 0.3678 | 0.3334 |
| GEX Reads mapped confidently to transcriptome | 0.6146 | 0.6536 |
| GEX Reads mapped to genome | 0.9151 | 0.9261 |
| GEX Reads with TSO | 0.3042 | 0.3386 |
| GEX Sequenced read pairs | 2593356890 | 870279531 |
| GEX Total genes detected | 28570 | 28398 |
| GEX Valid UMIs | 0.9995 | 0.9739 |
| GEX Valid barcodes | 0.9363 | 0.829 |

**Supplementary Table 2 – List of genes considered transcription factors (TFs) for SCENIC+ analysis**

| | | | | | | | |
|---|---|---|---|---|---|---|---|
| Bcl6b | Zscan26 | Mtf1 | Klf9 | Zic5 | Zfp410 | Zfp3 | Zfp691 |
| Zfp637 | Egr3 | Klf12 | Bcl6 | Tfap2a | Tfap2b | Tfap2c | Tfap2e |
| Arid3a | Arid5a | Ascl2 | Tcf3 | Bhlhe40 | Myf6 | Max | Mafk |
| Atf1 | Jdp2 | Mafb | Glis2 | Plagl1 | Osr2 | Sp4 | Klf7 |
| Zbtb7b | Zic1 | Egr1 | Zfp281 | Hic1 | Zfp740 | Osr1 | Zbtb14 |
| Zbtb12 | Zscan4c | Zfp105 | Zfp128 | Zic2 | Zic3 | Zbtb3 | E2f3 |
| E2f2 | Spi1 | Elf3 | Gabpa | Ehf | Spdef | Foxj3 | Foxj1 |
| Foxa2 | Foxk1 | Foxl1 | Gata6 | Gata3 | Gata5 | Gcm1 | Six6 |
| Nkx3-1 | Hnf1a | Hoxa3 | Irf9 | Irf3 | Irf4 | Irf6 | Irf5 |
| Srf | Myb | Mybl1 | Rxra | Hnf4a | Esrra | Nr2f2 | Rara |
| Rfx4 | Rfx7 | Rfx3 | Sp100 | Gmeb1 | Smad3 | Tcf7 | Hbp1 |
| Cic | Bbx | Sox8 | Tcf7l2 | Sox17 | Lef1 | Sox30 | Sox15 |
| Sox5 | Sox18 | Sox12 | Sox14 | Tcf7l1 | Sox21 | Sox7 | Sox11 |
| Sry | Sox13 | Sox4 | Sox1 | Eomes | Tbp | Cux1 | Lhx2 |
| Hoxb6 | Hoxa10 | Hoxa4 | Meox1 | Nkx2-1 | Irx2 | Dlx3 | Mnx1 |
| Hoxc13 | Hoxc11 | Hoxc8 | Hoxc6 | Evx2 | Hoxd13 | Evx1 | Otx1 |
| Rhox6 | Vax1 | Pknox1 | Phox2a | Phox2b | Alx3 | Hoxa2 | Nkx2-5 |
| Lhx1 | Lhx9 | Meis1 | Hnf1b | Dlx4 | Hoxb9 | Gsc | Barx1 |
| Msx2 | Pitx1 | Irx4 | Otp | Otx2 | Hmbox1 | Hoxc10 | Hoxc5 |
| Dlx2 | Esx1 | Six2 | Rax | Cdx1 | Pitx3 | Msx3 | Lhx4 |
| Prrx1 | Lmx1a | Barhl1 | Lhx6 | Lhx3 | Hoxd8 | Meis2 | Nkx2-2 |
| Shox2 | Pitx2 | Dmbx1 | Nkx1-1 | Uncx | Lhx5 | Pdx1 | Cdx2 |
| Nobox | Dlx5 | Hoxa1 | Dbx1 | Isx | Irx3 | Irx5 | Irx6 |
| Barx2 | Isl2 | Vsx1 | Barhl2 | Six4 | Gbx2 | Hdx | Vax2 |
| Lbx2 | Nkx6-1 | Arx | Pknox2 | Gsx2 | Hoxc9 | Alx1 | Hoxa13 |
| Hoxa11 | Hoxa9 | Hoxa7 | Hoxa5 | Hoxb4 | Hoxb5 | Hoxb7 | Lmx1b |
| Six3 | En2 | Hlx | Prrx2 | Alx4 | Meis3 | Crx | Obox6 |
| Dlx1 | Hoxd1 | Hoxd11 | Hoxa6 | Emx2 | Nkx2-6 | Nkx2-3 | Prop1 |
| Dbx2 | Tgif1 | Msx1 | Duxbl1 | Nkx1-2 | Hoxb3 | Hoxb13 | Nkx3-2 |
| Hmx2 | Hoxc12 | Hoxd10 | Rhox11 | Six1 | Pbx1 | Nkx2-4 | Obox1 |
| Bsx | Hoxb8 | Homez | En1 | Nkx2-9 | Tgif2 | Nkx6-3 | Obox3 |
| Hmx1 | Gbx1 | Tlx2 | Obox5 | Obox2 | Hoxc4 | Hoxd3 | Cphx1 |
| Lhx8 | Pax6 | Pax7 | Pax4 | Pou1f1 | Pou2f2 | Pou6f1 | Pou4f3 |
| Pou2f1 | Pou2f3 | Pou3f4 | Pou3f1 | Pou3f2 | Pou3f3 | Ahctf1 | Hmga2 |
| Mlx | Srebf1 | Tfec | Nhlh2 | Xbp1 | Atf3 | Atf4 | Junb |
| Cebpb | Dbp | Zscan20 | Sp1 | Zfp263 | Zscan10 | Zfp202 | Zkscan1 |
| Mzf1 | Zfp300 | Zbtb1 | Egr2 | Klf8 | Snai1 | Dnajc21 | Zkscan5 |
| Zfx | Dmrtc2 | Foxj2 | Foxg1 | Foxp2 | Foxp1 | Foxo4 | Foxo1 |
| Foxc2 | Foxo3 | Foxo6 | Gata4 | Irf2 | Mecp2 | Mybl2 | Mypop |
| Prdm11 | Rarg | Nr2f6 | Nr2c1 | Esr1 | Nr2e1 | Esrrb | Nr5a2 |
| Esrrg | Nr4a2 | Rora | Rorb | Ar | Nr2f1 | Gmeb2 | Sox10 |
| Sox3 | Sox6 | Tbx2 | Tbx4 | Tbx1 | Tbx5 | Tbx3 | Tbx20 |
| Creb1 | Cebpa | Jun | Cebpg | Jund | Foxm1 | Foxn1 | Foxn4 |
| Gli3 | Gli1 | Gli2 | Tbr1 | Esr2 | Runx2 | Etv3 | Etv1 |
| Spic | Elk3 | Elk1 | Etv5 | Fli1 | Etv4 | Elk4 | Elf5 |

| | | | | | | | |
|---|---|---|---|---|---|---|---|
| Etv6 | Elf4 | Ets1 | Elf2 | Erg | Arid3b | Arid5b | Setbp1 |
| Phf21a | Tfe3 | Mnt | Tfap4 | Twist2 | Arnt | Arnt2 | Hey2 |
| Srebf2 | Hes1 | Hes7 | Tfeb | Bhlhe22 | Msc | Npas2 | Usf1 |
| Sohlh2 | Hes2 | Clock | Figla | Bhlhe41 | Tcf12 | Mitf | Mlxip |
| Tcfl5 | Olig3 | Tcf21 | Hes5 | Neurog1 | Bhlha15 | Tcf4 | Arntl |
| Usf2 | Atoh1 | Tef | Fosl1 | Atf2 | Fosl2 | Creb3l2 | Crebzf |
| Mafg | Cebpe | Nfil3 | Crem | Atf7 | Zfp711 | Foxc1 | Foxb1 |
| Foxl3 | Sebox | Nanog | Noto | Irf7 | Terf2 | Myrf | Rxrg |
| Rarb | Rxrb | Sp110 | Hmg20b | Mecom | Pax2 | Nr2e3 | Spz1 |
| Hlf | Creb5 | Zfp652 | Meox2 | Hoxd9 | Vdr | Trp73 | Trp53 |
| Rfx2 | Zeb1 | Pparg | Nr3c1 | Smad4 | Klf4 | Ctcf | Tfcp2l1 |
| Pou5f1 | Rela | Sox2 | Stat3 | Myod1 | Myog | Nfe2l2 | Gfi1b |
| Klf1 | Gata1 | Rfx1 | Runx1 | Stat6 | Stat4 | Ahr | Ascl1 |
| Hif1a | Myc | Mxi1 | Ptf1a | Neurog2 | Tal1 | Lyl1 | Neurod1 |
| Twist1 | Mycn | Hand1 | Neurod2 | Olig2 | Msgn1 | Nhlh1 | Fosb |
| Fos | Ddit3 | Bach1 | Batf3 | Batf | Nfe2l1 | Bach2 | Maff |
| Maf | Nfe2 | Klf6 | Klf5 | Zbtb17 | Wt1 | Ikzf1 | Sp2 |
| Yy1 | Snai2 | Ovol1 | Sp3 | Sall4 | Klf3 | Rest | Gfi1 |
| Prdm5 | Klf15 | Maz | Hinfp | Zbtb7a | Zfp57 | Ovol2 | Prdm1 |
| Prdm16 | Zfp335 | Prdm14 | Zfp322a | Zbtb33 | Zfp42 | Prdm9 | Sp7 |
| Zfp143 | Zfp932 | Insm1 | Ctcfl | Sp5 | Nfya | Rbpj | Cux2 |
| Onecut1 | Dmrt1 | Dmrtb1 | E2f4 | E2f7 | E2f1 | E2f5 | E2f6 |
| Ebf1 | Etv2 | Spib | Ets2 | Elf1 | Fev | Foxa1 | Foxq1 |
| Foxa3 | Foxi1 | Foxl2 | Foxd3 | Gata2 | Grhl2 | Vsx2 | Cdx4 |
| Pbx2 | Pbx3 | Isl1 | Hsf2 | Hsf1 | Irf1 | Irf8 | Mef2d |
| Mef2c | Mef2a | Mbd2 | Nr1h3 | Nr1i3 | Nr2c2 | Hnf4g | Nr1d1 |
| Nr1d2 | Ppara | Nr1i2 | Nr4a1 | Nr5a1 | Rorc | Pgr | Nr1h4 |
| Thra | Trp63 | Pax5 | Relb | Rel | Nfatc4 | Nfkb2 | Nfatc2 |
| Nfkb1 | Nfatc3 | Nfatc1 | Rfx6 | Runx3 | Aire | Nfib | Nfia |
| Nfic | Sox9 | Stat5a | Stat5b | Stat1 | Stat2 | Tbx21 | T |
| Tead4 | Tead2 | Tead1 | Thap11 | Nfyb | Epas1 | Taf1 | Cbfb |
| Nfyc | Nrf1 | Myf5 | Mafa | Rbpjl | Nr1h2 | Hif3a | Creb3 |
| Nfe2l3 | Peg3 | Gtf3a | E4f1 | Bnc1 | Sall1 | Klf16 | Ikzf5 |
| Zfp697 | Plagl2 | Zfp182 | Zfp1 | Zfp467 | Zfp664 | Ebf2 | Ebf4 |
| Foxd1 | Mixl1 | Mef2b | Smad5 | Smad9 | Fank1 | Zfp384 | Tfap2d |
| Brca1 | Cdc5l | Cebpd | Cxxc1 | Egr4 | Foxf1 | Foxf2 | Foxp3 |
| Fubp1 | Glis3 | Hesx1 | Hltf | Hmga1 | Hmga1b | Hoxb1 | Hoxd4 |
| Mlxipl | Nfat5 | Nr0b1 | Nr4a3 | Nr6a1 | Onecut2 | Pax3 | Pax8 |
| Plag1 | Pou4f2 | Ppard | Pura | Rreb1 | Smad1 | Smad2 | Smarca5 |
| Tead3 | Tfcp2 | Tfdp1 | Thrb | Ubp1 | Zbtb18 | Zbtb6 | Hivep2 |
| Zfp148 | Zfp423 | Dmc1 | Il21 | Tbx6 | Zfp809 | Frem1 | Prdm15 |
| Hmx3 | Prdm4 | Bcl11b | Ikzf3 | Npas4 | Atf5 | Atf6 | Bcl11a |
| Bcl3 | Bhlhe23 | Bptf | Cebpz | Cenpb | Chd1 | Chd2 | Cpeb1 |
| Creb3l1 | Dlx6 | E2f8 | Ebf3 | Emx1 | Ep300 | Erf | Ezh2 |
| Foxd2 | Gcm2 | Glis1 | Grhl1 | Gsc2 | Gsx1 | Gtf2f1 | Gtf2i |
| Gzf1 | Hcfc1 | Hey1 | Hic2 | Hivep1 | Hoxb2 | Hoxd12 | Hsf4 |
| Hsfy2 | Id4 | Ikzf2 | Klf13 | Klf14 | Mesp1 | Mga | Mta3 |
| Nfix | Nkx6-2 | Nr3c2 | Nrl | Olig1 | Onecut3 | Patz1 | Pax1 |

| | | | | | | | |
|---|---|---|---|---|---|---|---|
| Pax9 | Pml | Pou4f1 | Pou6f2 | Prox1 | Prrxl1 | Rad21 | Rcor1 |
| Rfx5 | Rhox4e | Scrt1 | Scrt2 | Sin3a | Six5 | Smarcc1 | Smarcc2 |
| Smc3 | Sp8 | Tbl1xr1 | Tbx15 | Tbx19 | Tgif2lx1 | Tgif2lx2 | Tlx1 |
| Wrnip1 | Ybx1 | Yy2 | Zbtb16 | Zbtb4 | Zbtb49 | Zbtb7c | Zfhx3 |
| Zfp110 | Zfp219 | Zfp282 | Zfp524 | Zfp784 | Zic4 | Zkscan3 | Gabpb1 |
| Rfxap | Gtf2a2 | Pou2af1 | Foxh1 | Smad6 | Smad7 | Rfxank | Deaf1 |
| Tal2 | Mxd3 | Hand2 | Zfp628 | Dmrt2 | Dmrt3 | Dmrta1 | Dmrta2 |
| Zfp354c | Tbx22 | Trim28 | Gtf2ird1 | Cnot3 | Mbtps2 | Satb1 | Gtf2b |
| Klf11 | Zscan4f | Zscan4d | Ctnnb1 | Klf10 | Lbx1 | Zfp692 | Zfp800 |
| Foxd4 | Zfp653 | Klf2 | Zfp276 | Zfp445 | Adnp2 | Klf17 | Tbx10 |
| Mynn | Insm2 | Zfp414 | Zfp654 | Sp6 | Snai3 | Foxe3 | Zfp575 |
| Aebp2 | Zfp661 | Zfp641 | Fezf2 | Zfp526 | Zbtb5 | Zscan12 | Zfp536 |
| Ikzf4 | Snapc4 | Zfp579 | Zfp513 | Zfp91 | Zfp426 | Zfp92 | Zkscan6 |
| Zfp2 | Zfy1 | Zscan21 | Fezf1 | Zfp41 | Zfp239 | Zfy2 | Zfp367 |
| Zfp710 | Zfp553 | Zfp74 | Zfp775 | Zbtb24 | Zfp574 | Zik1 | Zbtb41 |
| Zfp184 | Zfp768 | Zfp24 | Zfp319 | Zfp64 | Zfp672 | Zkscan14 | Zfp37 |
| Zfp39 | Zscan2 | Zfp112 | Zfp354a | Zfp667 | Zfp583 | Zfp260 | Zfp329 |
| Zfp629 | Zfp787 | Zfp647 | Zbtb20 | Zfp786 | Zfp689 | Zfp451 | Zfp668 |
| Zfp771 | Zfp46 | Zfp770 | Zfp523 | Zfp287 | Zfp422 | Zfp580 | Zfp354b |
| Zfpm1 | Zfp507 | Zbtb22 | Sall2 | Zfp280d | Zfp35 | Vezf1 | Zfp592 |
| Zfp382 | Zfp532 | Zfp516 | Sall3 | Zfp93 | Zfp90 | Zfp131 | Zfp82 |
| Hdac1 | Ctbp1 | Myef2 | Gm9833 | Carf | Nono | Id2 | Arntl2 |
| Helt | Zeb2 | Lmo2 | Ldb1 | Gtf2a1 | Gm28047 | Ilf2 | Taf6 |
| Mxd1 | Mxd4 | Mycs | Apex1 | Gm9044 | Gm32802 | Gm9049 | Gm9040 |
| Gm32717 | Gm9045 | Gm9048 | Gm9046 | Pbx4 | Zfhx2 | Atf6b | Thap1 |
| Drap1 | Foxk2 | Gatad1 | Gatad2a | Hmg20a | Hmgxb4 | Kmt2b | Lcorl |
| Mbd1 | Ncoa1 | Phf20 | Ski | Zbtb11 | Zbtb21 | Zbtb25 | Zbtb2 |
| Zbtb40 | Zgpat | Zfp217 | Zfp362 | Zfp444 | Zfp511 | Zfp518a | Zfp639 |
| Zfp644 | Rhox10 | Zfp341 | Trps1 | Nf1 | Atoh7 | Creb3l4 | Ferd3l |
| Foxe1 | Sp9 | Tbx18 | Bnc2 | Zfp180 | Zfp65 | Zfp398 | Zkscan17 |
| Zfp764 | E430018J 23Rik | Cpsf4 | Foxn2 | Gm28230 | Mesp2 | Skor1 | Zfp597 |
| Ing4 | Churc1 | Ltf | Rb1 | Foxr1 | Bclaf1 | Ccnt2 | Setdb1 |
| Hdac2 | Zfp366 | Dpf2 | Ahdc1 | Tfdp2 | Foxp4 | Hmgxb3 | Paxip1 |
| Satb2 | Gpbp1l1 | Zbtb44 | Zfp383 | Zfp369 | Gm44973 | Zfp655 | Phf1 |
| Mtf2 | Zscan29 | Cxxc5 | Foxn3 | Msantd3 | Zzz3 | Kdm2b | Kmt2a |
| Dnmt1 | Tet1 | Zhx1 | Rhox13 | Crebl2 | Foxb2 | Nfx1 | Ash2l |
| Mettl14 | Vps72 | Hes6 | Dpf1 | Zbtb32 | Zfp821 | Zfp174 | Zbtb45 |
| Zscan5b | Zfp12 | Zfp704 | Zfp612 | Zbtb43 | Zfp296 | Zbtb26 | Zfp449 |
| Zfp454 | Zbtb37 | Zfp113 | Zfp11 | Mbnl2 | Lin28b | Foxr2 | Tlx3 |
| Irx1 | Hsf5 | Skor2 | Xpa | Zfp14 | Zkscan7 | Zfp777 | Gm49345 |
| Zfp595 | Zfp953 | Zfp738 | Zfp457 | Hhex | Cers4 | Cers2 | Cers5 |
| Cers6 | Cers3 | Zkscan2 | Gm3854 | Zfp324 | Zfp13 | Zfp94 | Zfp558 |
| Zfp78 | Zfp317 | Zfp189 | Rbak | Zfp7 | Gm49527 | Zfp763 | Zfp212 |
| Zkscan16 | Zfp791 | Zfp266 | Zfp941 | 2610021A 01Rik | Gm28360 | Gm28168 | Zfp54 |
| Gm7145 | Zfp9 | BC02592 0 | Zfp169 | Zfp961 | Zfp566 | Gm26920 | Zfp69 |

| | | | | | | | |
|---|---|---|---|---|---|---|---|
| Zfp605 | Zfp429 | Gm28557 | Zfp874a | Zfp87 | Zfp85 | Zfp493 | Zfp72 |
| Zfp874b | Zfp273 | Rsl1 | Zfp456 | Zfp455 | Rslcan18 | Zfp58 | Zfp708 |
| Zfp607b | Zfp59 | Zfp60 | Zfp850 | Zfp607a | Zfp626 | Zfp248 | Zfp334 |
| Zfp872 | Gm20422 | Zfp879 | Zscan22 | Prdm6 | Zfp213 | Zfp146 | Zbtb42 |
| Zfp985 | Zfp989 | Zfp978 | Zfp992 | Zfp980 | Zfp760 | Zfp993 | Zfp534 |
| Zfp943 | Zfp987 | Zfp981 | Zfp995 | Gm21411 | Zfp994 | Zfp947 | Zfp991 |
| Zfp944 | Gm7072 | Zfp946 | Rex2 | Zfp986 | Zfp820 | Zfp990 | Zfp942 |
| Zfp988 | Zfp984 | Zfp945 | Zfp979 | Zfp28 | Zfp560 | Zbtb48 | Zfp541 |
| Prdm12 | Zfp236 | Zfp521 | Zfp316 | Mterf1b | Mterf1a | Dnttip1 | Topors |
| Adnp | Champ1 | Dach1 | Kat7 | Ncoa3 | Nfxl1 | Prdm10 | Zbtb10 |
| Zbtb8a | Gm44505 | Zkscan8 | Zfp407 | Zfp512 | Zfp84 | Aff4 | Ascc1 |
| Bad | Cbfa2t2 | Zfp830 | Cnot6 | Nelfb | Ddx20 | Eno1b | Eno1 |
| Fhl2 | Gtf2h3 | Gtf3c2 | Gm29609 | Gtf3c5 | Hcfc2 | Hcls1 | Hdac8 |
| Ube2k | Htatip2 | Kdm5a | Larp1 | Ncald | Nme1 | Nmral1 | Nucb1 |
| Otud4 | Pdcd11 | Pdlim5 | Phtf1 | Pir | Pqbp1 | Purg | Rab18 |
| Rbbp5 | Rbfox2 | Scmh1 | Sema4a | Sf1 | Snapc5 | Snd1 | Ssbp3 |
| Gm5751 | Ssxa1 | Ssxb2 | Gm6592 | Ssxb6 | Ssxb1 | Ssx9 | Gm14459 |
| Ssxb3 | Ssxb8 | Ssxb10 | Gm21876 | Ssxb5 | Ssxb9 | Taf1a | Taf9 |
| Ak6 | Tbpl1 | Tfam | Med30 | Timeless | Trmt1 | Tsc22d4 | Tulp1 |
| Vps4b | Yeats4 | Zbtb46 | Zhx3 | Zfp160 | Zfp207 | Rnf114 | Zfp326 |
| Zfp385a | Zfp503 | Zfp706 | Smarca1 | Zfp622 | Tbpl2 | Traf4 | Tcf15 |
| Sirt6 | Parp1 | Terf1 | Brf1 | Bdp1 | Polr3a | Ewsr1 | Tmem33 |
| Acaa1a | Acaa1b | Ovol3 | Zfp688 | Rlf | Zfp746 | Zfp687 | Zfp438 |
| Zscan18 | Prdm13 | Gtf2a1l | Zbtb8b | Zfp280b | Zbtb34 | Zscan25 | Fiz1 |
| Zfp853 | Zxdc | Zfp648 | Zfp408 | Zfp871 | Zfp882 | Zfp563 | Zfp952 |
| Zfp955b | Zfp870 | Zfp81 | Zfp617 | Zfp955a | Zfp709 | Zfp799 | Zfp811 |
| Zfp472 | Zfp101 | Zfp286 | Zfp30 | Zfp109 | Zfp114 | Zfp108 | Zfp235 |
| Zfp623 | Zfp551 | Zfp62 | Gm4924 | Zfp397 | Zfp790 | Zfp780b | Zfp729b |
| Zfp459 | Zfp458 | Zfp748 | Zfp759 | Zfp729a | Zfp712 | Zfp606 | Zkscan4 |
| Zfp251 | Zfp358 | Zfp61 | Zfp651 | Zbtb39 | Zfp646 | Zbtb38 | Banp |
| Crtc2 | Sfpq | Abl1 | Dido1 | Hnrnpul1 | Ncoa2 | Ilf3 | Lcor |
| Mllt10 | Zfp142 | Zfp462 | Yod1 | Mettl3 | 1700009N14Rik | 1700080O16Rik | 1810024B03Rik |
| 2010315B03Rik | 2310011J03Rik | 2410141K09Rik | 2610044O15Rik8 | 2810403A07Rik | 3300002I08Rik | 3830417A13Rik | 4921501E09Rik |
| 4921509C19Rik | 5730507C01Rik | 6720489N17Rik | 9030624G23Rik | 9130019O22Rik | 9130023H24Rik | A1cf | Abcf2 |
| Aco1 | Adarb1 | Agap2 | Aggf1 | Agmat | Ahrr | AI987944 | Akr1a1 |
| Anxa1 | Anxa11 | Apex2 | Arfgap1 | Arg1 | Arg2 | Arid3c | Asap3 |
| Aspscr1 | Atoh8 | AU041133 | Aven | AW146154 | AW822073 | B230307C23Rik | Bax |
| BC005561 | Bmyc | Boll | Borcs8 | Brf2 | Canx | Cat | Cbx3 |
| Cbx7 | Ccdc25 | Cd59a | Cd59b | Cdk2ap1 | Celf4 | Celf5 | Celf6 |
| Cfl2 | Ckmt1 | Clk1 | Cnot4 | Cptp | Csnk2b | Cstf2 | Ctbp2 |
| Cxx1a | Cxx1b | Cxx1c | Cyb5r1 | Cycs | D130040H23Rik | D3Ertd254e | Dab2 |
| Dazap1 | Ddx4 | Ddx43 | Dgcr8 | Dhx36 | Diablo | Dis3 | Dmap1 |

| | | | | | | | |
|---|---|---|---|---|---|---|---|
| Dnmt3a | Dr1 | Dtl | Dus3l | Dusp22 | Dusp26 | Duxbl2 | Duxbl3 |
| Duxf3 | Ecsit | Edn1 | Eef1akmt3 | Eef1d | Eif5a2 | Esrp1 | Esrp2 |
| Etfb | Exo5 | Exosc3 | Ezr | Faap24 | Fbxl19 | Fez1 | Fgf15 |
| Foxi2 | Foxi3 | Foxs1 | Gadd45a | Gar1 | Git2 | Glyctk | Gm10053 |
| Gm10093 | Gm10130 | Gm10269 | Gm10324 | Gm10479 | Gm10662 | Gm10668 | Gm10770 |
| Gm10778 | Gm11007 | Gm12166 | Gm12184 | Gm12845 | Gm13212 | Gm14147 | Gm14151 |
| Gm14288 | Gm14295 | Gm14305 | Gm14308 | Gm14322 | Gm14325 | Gm14326 | Gm14327 |
| Gm14391 | Gm14399 | Gm14403 | Gm14406 | Gm14410 | Gm14412 | Gm14418 | Gm14419 |
| Gm14434 | Gm14435 | Gm14440 | Gm14443 | Gm14444 | Gm15446 | Gm17655 | Gm2000 |
| Gm2004 | Gm2007 | Gm2026 | Gm20939 | Gm28308 | Gm3055 | Gm35315 | Gm3604 |
| Gm38394 | Gm43517 | Gm4513 | Gm45233 | Gm4565 | Gm4567 | Gm45871 | Gm4631 |
| Gm4724 | Gm4767 | Gm4881 | Gm4922 | Gm5157 | Gm5294 | Gm5454 | Gm5890 |
| Gm5891 | Gm6176 | Gm6710 | Gm6811 | Gm6871 | Gm6882 | Gm6902 | Gm7168 |
| Gm7356 | Gm8444 | Got1 | Gpam | Gpank1 | Gpd1 | Grhl3 | Grhpr |
| Gtpbp1 | Gtpbp6 | H1fx | H2afy | H2afz | Hdac3 | Hdac6 | Heyl |
| Hhat | Hirip3 | Hist2h2ab | Hivep3 | Hlcs | Hmga1-rs1 | Hmgb1 | Hmgb2 |
| Hmgb3 | Hmgb4 | Hmgn3 | Hnrnpa0 | Hnrnpc | Hnrnph3 | Hnrnpll | Hp1bp3 |
| Hsf3 | Hspa1l | Hspa5 | Id1 | Il24 | Ing3 | Ivd | Jazf1 |
| Kat2a | Kcnip1 | Kdm2a | Kdm4a | Kdm4b | Kdm4c | Kdm4d | Kdm5b |
| Kdm5d | Kdm7a | Kif22 | Klrg1 | Larp4 | Las1l | Lin28a | Lrrfip1 |
| Lsm6 | Luzp1 | Luzp2 | Magea1 | Magea10 | Magea2 | Magea3 | Magea4 |
| Magea5 | Magea6 | Magea8 | Magoh | Map4k2 | Mapk1 | Mbd4 | Mctp2 |
| Mdm2 | Mex3c | Mief1 | Mios | Mkx | Morn1 | Mrpl1 | Mrpl2 |
| Mrps25 | Msi1 | Msi2 | Msra | Msrb3 | Mthfd1 | Mycl | Mylk |
| Nags | Nanos1 | Nap1l1 | Ncbp2 | Ncor1 | Ncor2 | Nelfa | Nelfe |
| Neurog3 | Nmi | Nnt | Noc2l | Npdc1 | Nr1h5 | Nuak1 | Nuak2 |
| Nup107 | Nup133 | Nxph3 | Obox3-ps8 | Odc1 | P4hb | Pck2 | Pde6h |
| Pds5a | Pgam2 | Phf2 | Phf8 | Phlda2 | Pick1 | Pik3c3 | Pkm |
| Plg | Pold2 | Pole3 | Pole4 | Poli | Polr3g | Ppargc1a | Ppp1r10 |
| Ppp2r3d | Ppp5c | Prdx5 | Prkaa1 | Prkaa2 | Prnp | Prox2 | Psma6 |
| Psmc2 | Psmd12 | Ptcd1 | Pum3 | R3hdm2 | Rab14 | Rab2a | Rab7 |
| Ran | Rbbp9 | Rbm17 | Rbm22 | Rbm3 | Rbm42 | Rbm7 | Rbm8a2 |
| Rbms1 | Rfc2 | Rfc3 | Rfx8 | Rhox12 | Rnaseh2c | Rnf138 | Rpl35 |
| Rpl6 | Rpp25 | Rps10 | Rps4x | Rps6ka5 | Rufy3 | Ruvbl1 | Sap30 |
| Scx | Sf3b1 | Sft2d1 | Sim1 | Sim2 | Slc18a1 | Smap2 | Smarca4 |
| Smarcb1 | Smok2a | Smok2b | Smok3a | Smok3b | Smok3c | Smpx | Smug1 |
| Snrnp70 | Snrpb2 | Socs4 | Sod1 | Spag7 | Spats2 | Spr | Srbd1 |
| Srp9 | Srrm3 | Ssrp1 | Stau2 | Stub1 | Suclg1 | Supt20 | Suz12 |
| Taf7 | Tagln2 | Tceal3 | Tceal5 | Tceal6 | Tcf23 | Tcf24 | Tff3 |
| Thap12 | Thoc2 | Tia1 | Timm44 | Timm8a1 | Timm8a2 | Tob2 | Tpi1 |
| Tppp | Trerf1 | Trim21 | Trim24 | Trim33 | Trim66 | Trim69 | Trip10 |
| Trmo | Trove2 | Tsn | Tsnax | U2af1 | Ube2v1 | Ubtf | Ubxn1 |
| Ugp2 | Uqcrb | Usp39 | Utp18 | Vamp3 | Wdr83 | Wisp2 | Xrcc1 |
| Xrcc4 | Ywhae | Ywhaz | Zbed6 | Zc3h11a | Zc3h7a | Zcchc14 | Zcchc17 |
| Zdhhc15 | Zdhhc24 | Zdhhc5 | Zfa-ps | Zfp111 | Zfp119a | Zfp119b | Zfp120 |
| Zfp27 | Zfp275 | Zfp352 | Zfp386 | Zfp420 | Zfp433 | Zfp442 | Zfp568 |
| Zfp598 | Zfp683 | Zfp707 | Zfp719 | Zfp747 | Zfp781 | Zfp819 | Zfp825 |

| | | | | | | | |
|---|---|---|---|---|---|---|---|
| Zfp831 | Zfp846 | Zfp866 | Zfp867 | Zfp868 | Zfp869 | Zfp930 | Zfp931 |
| Zfp933 | Zfp935 | Zfp937 | Zfp938 | Zfp949 | Zfp950 | Zfp951 | Zfp958 |
| Zfp959 | Zfp960 | Zfp963 | Zfp964 | Zfp965 | Zfp966 | Zfp967 | Zfp968 |
| Zfp969 | Zfp97 | Zfp970 | Zfp971 | Zfp973 | Zfp974 | Zfp975 | Zfp976 |
| Zhx2 | Zmat2 | Zmat4 | Zmiz1 | Zrsr1 | Zscan4-ps1 | Zscan4-ps2 | Zscan4-ps3 |
| Zscan4b | Zscan4e | Zswim1 | Zxdb | | | | |
